# Single-Cell Epigenomics Uncovers Heterochromatin Instability and Transcription Factor Dysfunction during Mouse Brain Aging

**DOI:** 10.1101/2025.04.21.649585

**Authors:** Maria Luisa Amaral, Sainath Mamde, Michael Miller, Xiaomeng Hou, Jessica Arzavala, Julia Osteen, Nicholas D. Johnson, Elizabeth Walker Smoot, Qian Yang, Emily Eisner, Qiurui Zeng, Cindy Tatiana Báez-Becerra, Jacqueline Olness, Joseph Colin Kern, Jonathan Rink, Ariana Barcoma, Silvia Cho, Stella Cao, Nora Emerson, Jasper Lee, Jackson Willier, Timothy Loe, Henry Jiao, Songpeng Zu, Quan Zhu, Allen Wang, Joseph R. Ecker, Maria Margarita Behrens, Bing Ren

## Abstract

The mechanisms regulating transcriptional changes in brain aging remain poorly understood. Here, we use single-cell epigenomics to profile chromatin accessibility and gene expression across eight brain regions in the mouse brain at 2, 9, and 18 months of age. In addition to a significant decline in progenitor cell populations involved in neurogenesis and myelination, we observed widespread and concordant changes of transcription and chromatin accessibility during aging in glial and neuronal cell types. These alterations are accompanied by dysregulation of master transcription factors and a shift toward stress-responsive programs driven by AP-1, indicating a progressive loss of cell identity with aging. We also identify region- and cell-type-specific heterochromatin decay, characterized by increased accessibility at H3K9me3-marked domains, activation of transposable elements, and upregulation of long non-coding RNAs, particularly in glutamatergic neurons. Together, these results reveal age-related disruption of heterochromatin maintenance and transcriptional programs, identify vulnerable brain regions and cell types, and pinpoint key molecular pathways altered in brain aging.

**Highlights:**

- Single-cell multimodal profiling across eight brain regions reveals coordinated chromatin and transcriptional shifts during aging
- Age-related depletion of progenitor cells coincides with dysregulation of key developmental transcription factors
- Cell identity maintenance is compromised with the decline of master transcription factors
- Heterochromatin destabilization accompanied by activation of AP-1, transposable elements, pseudogene families, and long non-coding RNAs

## Introduction

Aging is accompanied by widespread cellular and molecular changes in the brain, which contribute to cognitive decline, reduced neural plasticity, and increased vulnerability to neurodegenerative diseases^1^. Detailed characterization of these changes holds the key to a better understanding of the mechanisms of organismal aging. While gene expression alterations have been extensively analyzed at single-cell resolution^2,3^, the transcriptional regulatory mechanisms shaping these changes remain largely unresolved. Epigenetic processes, manifested as chromatin accessibility, DNA methylation, histone modifications, and 3D chromatin organization, play a crucial role in defining cellular identity and responsiveness to environmental cues^4^, yet how epigenomic landscapes shift with age across different brain regions and cell types is not well understood. Here, we present a comprehensive analysis of cell-type- and brain-region-resolved chromatin accessibility of brain aging, integrating single-cell Assay for Transposase Accessible Chromatin using sequencing (ATAC-seq) and RNA-seq data across eight mouse brain regions. By resolving epigenomic changes at single-cell resolution, we uncover regulatory mechanisms that contribute to aging-associated transcriptional shifts and cellular dysfunction.

Previous studies have uncovered common hallmarks of aging—including genomic instability, stem cell exhaustion, mitochondrial dysfunction, and inflammation—shared across tissues and model organisms^5^. More recently, epigenetic alterations such as chromatin accessibility dynamics, DNA methylation changes, and histone modifications have emerged as key regulators of aging^6^, contributing to a progressive loss of transcriptional and regulatory fidelity that affects cellular identity and function^7^. Notably, epigenetic aging is also a reversible process, as ectopic expression of pluripotency factors^8^, activation of SIRT6 histone deacetylase^9,10^, and modulation of other histone-modifying enzymes^11,12^ have been shown to mitigate certain aging phenotypes. However, prior studies on brain aging largely rely on bulk tissue analyses, which obscure cell-intrinsic aging effects and fail to capture regulatory changes across different brain cell types and regions. Given the brain’s cellular diversity, a cell-type-resolved analysis of chromatin aging is essential to distinguish intrinsic regulatory changes from shifts in cellular composition, specially since neurodegenerative diseases disproportionately affect specific brain regions and cell populations—such as dopaminergic neurons in the substantia nigra in Parkinson’s disease^13^, oligodendrocytes in the white matter in multiple sclerosis^14^, and hippocampal neurons in Alzheimer’s disease^15^. A deeper understanding of how chromatin landscapes change during aging at the cell-type level will provide insights into how regulatory changes contribute to neurodegeneration and could uncover new therapeutic opportunities.

Previous studies have characterized transcriptomic changes in aging at the single-cell level, including large-scale efforts such as Tabula Muris Senis^2^, which cataloged aging signatures across multiple tissues, and a recent brain-wide atlas of aging^3^, which profiled 1.2 million neuronal and glial transcriptomes across the mouse brain. In addition to scRNA-seq, spatial transcriptomics has revealed white matter as particularly affected by aging^16^. These studies revealed that different cell types and spatial locations exhibit distinct transcriptional aging signatures. While transcriptomic atlases provide valuable insights, they alone do not capture the regulatory mechanisms driving transcriptional shifts.

The epigenome is essential for maintaining cell identity and gene regulation, yet how the epigenome changes with aging across different brain cell types and regions remains poorly understood. Focusing on chromatin accessibility, our previous study revealed that excitatory neurons in the mouse frontal cortex lose H3K9me3-marked heterochromatin with age, leading to increased accessibility at transposable elements and normally repressed genomic regions^17^. These findings suggest that heterochromatin instability may be a key feature of neuronal aging. However, a comprehensive, multi-region, single-cell chromatin accessibility map of the aging brain has been lacking. Here, we address this gap by profiling chromatin accessibility in 1 million cells across eight brain regions in both male and female mice. Additionally, our companion paper investigates age-associated DNA methylation changes and alterations in 3D genome organization.

Briefly, we report here a single-cell, multi-region chromatin accessibility and transcriptomic atlas of brain aging, generated through profiling of eight brain regions in male and female mice at 2, 9, and 18 months of age. Using a combination of single-cell assays including snATAC-seq, 10X multiome (single nucleus combined ATAC-seq and RNA-seq), and spatial transcriptomics, we examined the Prefrontal Cortex (FC), Anterior Hippocampus (AHC), Posterior Hippocampus (PHC), Entorhinal Cortex (EC), Nucleus Accumbens (NAC), Caudate Putamen (CP), Amygdala (AMY), and Periaqueductal Gray (PAG/PCG, RLP) (Table S1). These regions were selected due to their critical roles in cognition, memory, and emotional regulation, as well as their known vulnerabilities to age-related decline and neurodegenerative processes^18^. Integrative analysis showed a dramatic age-associated decline of three progenitor populations, accompanied by age-associated chromatin remodeling and transcriptional shifts. Major neuronal and glial populations display broad transcription factor dysregulation, chromatin accessibility changes, and disruptions in regulatory landscapes that drive shifts in cellular function. Finally, we observe widespread aging-associated chromatin changes, including heterochromatin destabilization, transposable element reactivation, and increased accessibility at stress-responsive elements, which emerge as defining features of brain aging. Together, this work provides insights into the transcriptional regulatory mechanisms underlying neuronal and glial aging in the mouse brain.

## Results

### Single-Nucleus Multi-Omics Analysis of the Aging Mouse Brains

To determine how chromatin accessibility and transcriptional programs shift during aging, we profiled eight brain regions across three ages (2, 9, and 18 months) in adult mice using single-nucleus ATAC-seq (snATAC-seq), single-nucleus Multiome (ATAC-seq + RNA-seq), and MERFISH spatial transcriptomics (Table S1). For snATAC-seq, we profiled each brain region in 2 biological replicates, each containing pooled brain tissues dissected from 3-7 mice, of 9-month-old and 18-month-old males, while incorporating our previously generated 2-month-old male data reported in^19^. For female samples, we generated single-nucleus 10X multiome data, extending our analysis to both modalities (ATAC-seq and RNA-seq) from all three age groups. In total, we analyzed chromatin accessibility in 1,116,638 cells, RNA expression in 368,491 cells, and spatial transcriptomics in 500,353 cells, obtaining a comprehensive view of age-dependent gene regulation and cellular composition (Figure 1a,b). We processed snATAC-seq data using SnapATAC2 and scRNA-seq + MERFISH data using Seurat^20^. Glial populations were broadly distributed across regions, while neuronal populations showed brain region specificity, consistent with region-specific neuronal specialization (Figure S1c). Batch correction with Harmony^21^ minimized technical effects between the male snATAC-seq and female 10x Multiome datasets, revealing no major batch effects (Figure S1b). Genome browser tracks of the age-associated gene C4b^22^ illustrate its consistent upregulation across brain regions, modalities, and sexes, demonstrating our ability to capture age-associated molecular changes (Figure S8a). More broadly, these changes in gene expression and chromatin accessibility are reflected as subtle gradients within cell clusters (Figure S1d-f). Among 80 distinct cell types identified using marker genes and integration with a single-cell transcriptomic atlas of the adult mouse brain^23^, the most abundant populations were oligodendrocytes, dentate gyrus (DG) glutamatergic neurons, and striatal D12 medium spiny neurons (Figure S1g). Notably, three progenitor cell populations showed significant age-related declines in cell abundance: Immature oligodendrocytes (IOL), OB-STR-CTX inhibitory immature neurons (IMN), and DG progenitors (Figure 1c,d).

**Figure 1.**
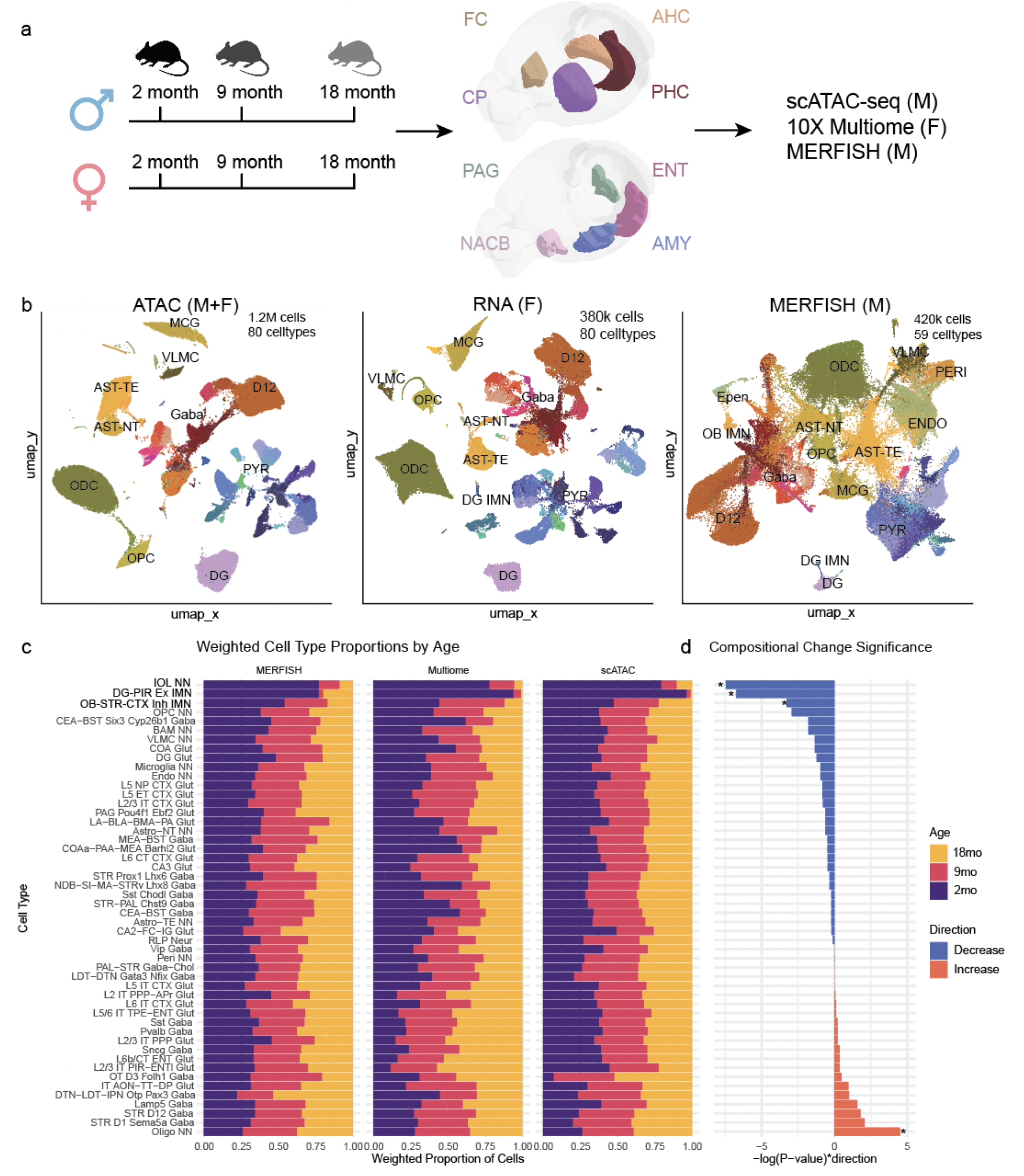
Single-nucleus multi-omic atlas of brain aging across eight regions in the mouse. (A) Schematic of experimental design showing brain regions profiled from male and female mice at 2, 9, and 18 months of age using snATAC-seq, 10X Multiome, and MERFISH. (B) Uniform Manifold Approximation and Projection (UMAP) of single-nucleus ATAC-seq (left), RNA-seq (center), and MERFISH (right) datasets, annotated by cell type. (C) Weighted cell-type proportions across ages for each modality, highlighting three progenitor populations that decline with age. (D) Statistical significance of age-associated changes in cell-type proportions (Wilcoxon rank-sum test).

To systematically characterize age-related chromatin remodeling, we identified open chromatin regions and candidate cis-regulatory elements (cCREs) in each cell type using MACS3^24^ (Methods), revealing 881,231 combined cCREs across all cell types (Figure S2a), with 724,636 of these overlapping a previous atlas of chromatin accessibility in the adult mouse brain^25^. A majority of cCREs were cell type-specific (Figure S2b). We assessed differential chromatin accessibility between 2-month-old and 18-month-old samples across three levels of resolution: All cell types combined, within each cell type across brain regions, and within each cell type further stratified by region and sex. We defined significant changes using an adjusted p-value < 0.01 and fold changes ( |logFC| > 0.5). This analysis revealed widespread chromatin remodeling with aging. At the most granular level, 88,555 cCREs exhibited significant increases in accessibility in at least one combination of celltype, region, and sex, with 30,994 unique to a single comparison. Conversely, 70,734 cCREs showed significant decreases, with 27,307 unique to a single combination. Some loci were recurrently altered across multiple comparisons: 492 cCREs were upregulated in ≥10 combinations of celltype, region, and sex, including a heterochromatin hotspot (chr13:68173417-68173918) previously linked to aging-related chromatin changes^17^. Strikingly, the most highly differential peaks across the genome were consistent between males and females, with several gene clusters—including the Killer cell lectin-like receptor (Klra), Secretoglobin (Scgb), Olfactory receptor (Olfr), Prostate and testis expressed (Pate), and Protocadherin gamma (Pcdhg) gene families—alongside intergenic regions, showing strong upregulation in neuronal aging (Figure S3a,b). Because the number of differentially accessible peaks correlates with cell number and complexity, the remainder of this study focuses on carefully characterizing cell type-specific chromatin remodeling. To complement our chromatin accessibility analysis, we performed differential gene expression analysis using MAST, adjusting for brain region, mitochondrial percentage, ribosomal percentage, and replicate as latent variables^26^. This analysis revealed widespread transcriptional changes across glutamatergic, GABAergic, and non-neuronal cell types with aging (Figure S4a, S5a, S6a; Methods). Aging alters chromatin accessibility and transcriptional programs in a cell-type-specific manner. In the following sections, we explore progenitor populations that decline with age and regulatory changes within major neuronal and glial cell types.

### Loss of Progenitor Populations in Aging Brains

Progenitor cells contribute to cellular renewal and plasticity during brain development. We identify three progenitor populations that experience dramatic declines in cell proportion with age. Among these, Dentate Gyrus (DG) progenitors exhibit the most striking reduction, decreasing from ~3% abundance in 2-month-old brains to <0.01% in both the anterior and posterior hippocampus (Figure 2d). These cells reside within one of the two major neurogenic niches in the adult brain—the dentate gyrus—where they give rise to excitatory granule neurons. The other principal neurogenic niche, the subventricular zone (SVZ), produces inhibitory neurons that migrate to the olfactory bulb. In line with this, we identified a second declining population—olfactory bulb immature inhibitory neuron-like cells annotated as OB-STR-CTX Inh IMN^23^. These cells are primarily found in the nucleus accumbens, caudate putamen, and hippocampus, comprising ~4% of cells at 2 months, but declining with age at varying rates (Figure 2d). The third progenitor population showing dramatic age-related decline is an intermediate cell type between oligodendrocyte progenitor cells and mature oligodendrocytes, known as immature oligodendrocytes (IOLs). This population consistently decreases across brain regions, from ~2% to ~0.1% with aging, aligning with prior findings of impaired remyelination in the aging brain^27^ (Figure 2c,d). These changes in progenitor cell proportions were observed in both the snATAC-seq and 10x Multiome datasets. Additionally, MERFISH spatial transcriptomic profiling confirmed the age-related, region-specific depletion of DG progenitors, OB-STR-CTX Inh IMN, and IOLs (Figure 2a–c). Notably, in aged brains, remaining progenitor cells were predominantly localized to canonical neurogenic niches. Because MERFISH captures intact brain slices rather than dissociated nuclei, it helps mitigate potential sampling biases inherent in single-nucleus profiling. Taken together, multiple lines of evidence indicate pronounced progenitor cell loss with aging.

**Figure 2.**
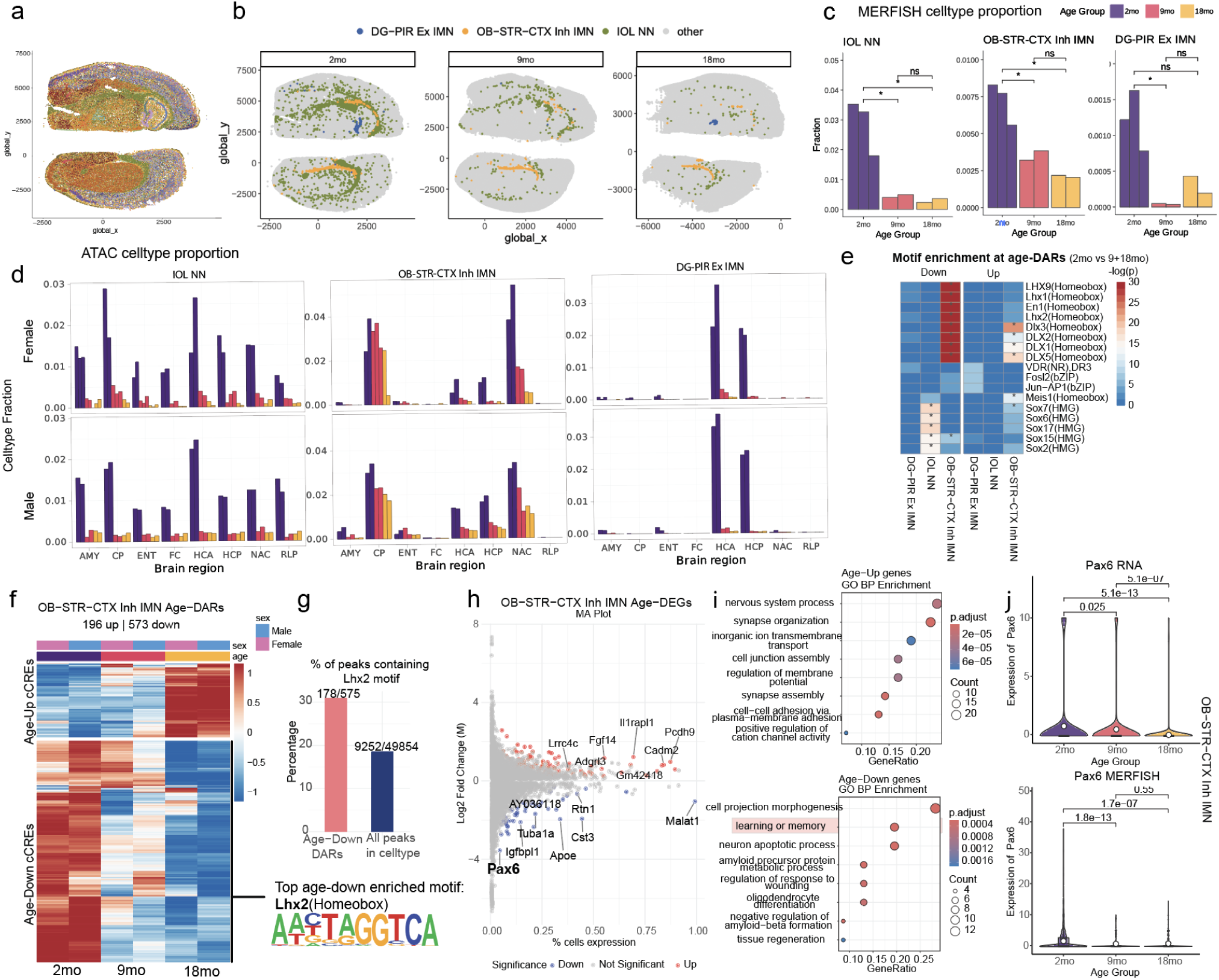
Aging depletes progenitor populations and disrupts neurogenic regulatory programs. (A) MERFISH spatial maps of brain slices (plates 46 and 69) with cells colored by annotated cell types (Table S1). (B) Representative MERFISH images across three ages showing immature oligodendrocytes (IOL), OB-STR-CTX inhibitory immature neurons (Inh IMN), and dentate gyrus (DG) progenitors. (C) Age-associated changes in progenitor cell-type proportions from MERFISH data. (D) Progenitor cell-type proportions from snATAC-seq across brain regions and sexes. (E) Motif enrichment analysis of age-associated differentially accessible cCREs in progenitor populations. (F) Heatmap showing normalized accessibility (z-score) of age-differential cCREs in OB-STR-CTX Inh IMN, stratified by age and sex. (G) Proportion of downregulated cCREs containing Lhx2 motifs in OB-STR-CTX Inh IMN; bottom shows the Lhx2 consensus motif. (H) MA plot of age-differential gene expression (18 mo vs. 2 mo) in OB-STR-CTX Inh IMN. (I) GO term enrichment for upregulated (top) and downregulated (bottom) genes in OB-STR-CTX Inh IMN. (J) Age-associated expression of *Pax6*, a direct target of Lhx2, in scRNA-seq (top) and MERFISH (bottom) datasets.

Recent studies suggest that progenitor cell exhaustion in aging is driven by chromatin dysfunction, including loss of repressive histone marks and altered chromatin accessibility, rather than a simple failure of proliferation^28^. By integrating chromatin accessibility and transcriptional datasets across multiple independent approaches, including MERFISH (male), snATAC-seq (male), and 10x Multiome (female), we confirm consistent reductions in progenitor cell populations across replicates. To determine whether chromatin changes accompany progenitor depletion, we analyzed age-differentially accessible cis-regulatory elements (cCREs) in progenitor cell populations across age groups. Motif enrichment analysis revealed a significant depletion of homeobox motifs in OB-STR-CTX Inh IMN and Sox motifs in IOL_NN cells, indicating dysregulation of cell-identity transcription factors (Figure 2e). We focus on OB-STR-CTX Inh IMN due to their abundance in our data and their lack of characterization in published literature. Differential accessibility analysis revealed a progressive shift in chromatin accessibility with aging, with 9-month-old samples exhibiting an intermediate profile between young (2-month) and aged (18-month) states, suggesting a gradual rather than abrupt epigenomic transition during aging (Figure 2f). Lhx2 emerged as the most significantly downregulated transcription factor in OB-STR-CTX Inh IMN, with age-associated decreases in accessibility at Lhx2-motif sites (Figure 2g). Lhx2 is a key transcriptional regulator of neurogenic progenitors, maintaining their self-renewal capacity and delaying differentiation into mature neuronal or glial fates^29,30^. In embryonic and adult neural stem cells, Lhx2 prevents premature differentiation and supports ongoing neurogenesis, in part by directly activating downstream targets such as *Pax6*^29,31^. Consistent with this, differential gene expression analysis between 2-month-old and 18-month-old samples revealed that *Pax6*, a direct target of Lhx2 and a key regulator of neurogenic lineage specification^31^, was significantly downregulated in both scRNA-seq and MERFISH data (adj p-value < 0.001, ~11.7 fold reduction in expression) (Figure 2j). *Pax6* is crucial for neural fate specification, and its age-related depletion underscores the progressive collapse of neurogenic potential. GO pathway enrichment analysis of genes differentially expressed between 2-month-old and 18-month-old samples revealed that those associated with learning, memory, and cell projection morphogenesis were downregulated with age (Figure 2i). Conversely, genes related to synapse organization were significantly upregulated in these progenitors (Figure 2i), suggesting that a subset of cells may be prematurely differentiating. While progenitor depletion is a well-documented feature of aging, the upregulation of synaptic genes in the remaining OB-STR-CTX Inh IMN population may reflect a disruption in the balance between undifferentiated progenitors and early postmitotic neurons. This shift may be linked to the loss of Lhx2 and other homeobox transcription factors, which normally sustain the neurogenic program and suppress premature differentiation^30^. Together, these findings support a model in which age-related chromatin remodeling leads to the erosion of progenitor identity in OB-STR-CTX Inh IMN, in part through the progressive loss of Lhx2 and *Pax6* transcriptional regulation.

### Dysregulation of cell identity genes in glial cells during aging

Having established the dramatic decline in progenitor populations, we next examined how aging affects the chromatin landscape of terminally differentiated cell types throughout the eight brain regions. We observed widespread transcriptional and chromatin accessibility changes in non-neuronal cell types, including oligodendrocytes, astrocytes, and microglia, which likely impact their ability to maintain brain homeostasis and support neuronal function. Dysfunction in these glial populations has been linked to neurodegenerative processes^32^, underscoring the importance of understanding their regulatory changes with age.

To systematically examine chromatin accessibility changes in oligodendrocytes, we identified 1727 up-regulated and 2501 down-regulated differentially accessible candidate cis-regulatory elements (cCREs) across aging (Figure S4a). Chromatin accessibility changes in oligodendrocytes were consistent across brain regions and sexes and progressed gradually from 2 to 18 months (Figure 3a). Similarly to immature oligodendrocytes (IOLs), mature oligodendrocytes exhibit a decline in the activity of Sox transcription factors, which are crucial for maintaining oligodendrocyte identity (Figure 2e, 3b)^33^. Motif enrichment analysis revealed significant enrichment of Sox family and Creb5 motifs in downregulated cCREs, while KLF and SP family motifs were enriched in upregulated peaks, marking a shift in transcription factor activity (Figure 3b). These changes were further confirmed by aggregate plots of Klf1 and Sox6 motif presence in upregulated, downregulated, and non-changing peaks across brain regions (Figure 3c). Differential gene expression analysis (Figure 3d) revealed a downregulation of genes related to myelin sheath formation and energy production, while genes associated with synaptic organization were enriched in upregulated genes (Figure 3e). Previous studies have shown that oligodendrocytes express synaptic genes, which participate in axon-myelin communication^34^. In this context, the upregulation of synaptic organization genes in aging oligodendrocytes may reflect an adaptive response to maintain axon-glial interactions as neural circuits change with age. On the other hand, the downregulation of myelination related genes was evident across the 8 brain regions (Figure S8b,c). *Mal*, a gene that functions in recruitment of myelin protein in oligodendrocytes^35^, is found to be significantly downregulated in oligodendrocytes from aged mice in both MERFISH and RNA-seq analysis (Figure 3h).

**Figure 3.**
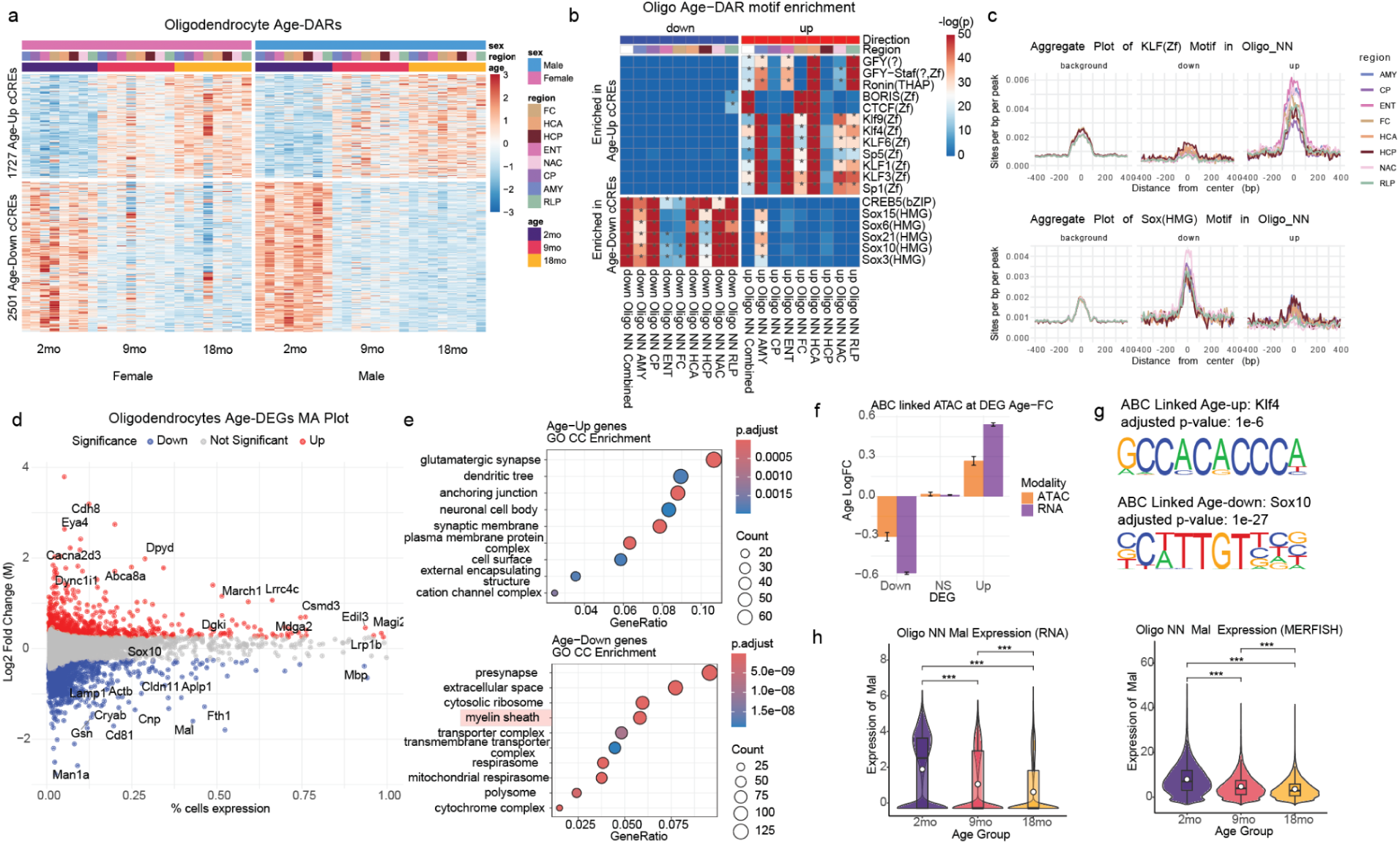
Aging oligodendrocytes exhibit transcriptional and chromatin changes associated with myelination loss and Sox factor decline. (A) Heatmap showing normalized accessibility (z-score) at age-differential cCREs in oligodendrocytes across brain regions and sexes. (B) Motif enrichment of downregulated (left) and upregulated (right) age-differential cCREs in oligodendrocytes; asterisks denote significance (adjusted *p* < 0.01). (C) Aggregate motif accessibility plots for Klf1 and Sox6 in upregulated, downregulated, and non-changing peaks in oligodendrocytes (Oligo NN). (D) MA plot of age-differential gene expression (18 mo vs. 2 mo) in oligodendrocytes. (E) GO term enrichment analysis for upregulated (top) and downregulated (bottom) genes. (F) Bar plot showing average age log fold-change of gene expression for downregulated, non-significant, and upregulated genes, with corresponding changes in linked peaks from ABC score analysis. (G) Motif logos for the most enriched motifs in peaks linked to upregulated (top) and downregulated (bottom) genes in aging oligodendrocytes. (H) Age-associated expression of the myelination gene *Mal* from scRNA-seq (left) and MERFISH (right).

We linked these differential genes to key regulatory cCREs in oligodendrocytes using Activity-By-Contact (ABC) analysis^36^, which predicts enhancer-promoter connections based on chromatin accessibility and genomic proximity. We calculated ABC scores using the snm3C-seq contact data from our companion paper and the accessibility data from our work in matching celltypes. We found that Sox10 was the most significantly enriched transcription factor motif within peaks linked to downregulated genes in aging oligodendrocytes (Figure 3g). Sox10 is a master regulator of oligodendrocyte lineage progression and myelin gene expression, playing a critical role in maintaining oligodendrocyte function and promoting myelination^33^. While Sox transcription factors are well-established as essential regulators of oligodendrocyte development, our findings reveal that their downregulation in aging could contribute to the well-documented loss of myelination in aged brains^37,38^. This suggests that, in addition to being critical for early oligodendrocyte differentiation, Sox factors are important in maintaining myelination throughout life, highlighting a previously underappreciated role for these transcription factors in aging-related myelin decline. Beyond transcription factor activity dysregulation and downregulation of important functions like myelination, aging oligodendrocytes exhibit increased expression of genes involved in inflammatory and regulatory pathways, including lncRNA *Neat1*, Parkinson’s disease driver gene *Snca*, and inflammatory marker *Il33* (Figure 3f). Collectively, these findings suggest that aging oligodendrocytes undergo chromatin and transcriptional changes that impair myelination, while the upregulation of synaptic and inflammatory genes, including *Snca* and *Il33*, may indicate a shift toward heightened responsiveness to neuronal and immune signals.

Astrocytes and microglia exhibit distinct shifts with aging, marked by increased accessibility at enhancers linked to stress-responsive transcription factors. Aging astrocytes from the frontal cortex exhibit the most age-dependent accessibility changes compared to other brain regions (Figure S4a). In these cells, cCREs gaining accessibility in aged mice are enriched for ATF(Zf) motifs, suggesting an adaptive response to the extracellular changes (Figure S4b). In contrast, cCREs gaining accessibility in aging microglia, particularly in microglia from the anterior hippocampus, exhibit IRF8 motif enrichment, indicating a shift toward a more pro-inflammatory, immune-activated state (Figure S4b). The upregulation of immune-related genes such as *Tnsfs8* and *Ildr2*, alongside the downregulation of classical microglial markers like C1qa and Cst3, suggests that aging microglia lose their cell identity while becoming chronically primed for inflammatory responses, a hallmark of neurodegeneration^1^. Previous studies have similarly reported that aging glial cells undergo transcriptional shifts marked by reduced activity of developmental transcription factors and increased expression of stress-responsive and immune-related genes^7,39–41^. Our brain-specific, multi-modal, and region-resolved analysis supports this model, revealing coordinated changes in chromatin accessibility and gene expression that underlie glial transcriptional reprogramming during aging.

### Upregulation of AP-1 Mediated Transcriptional Programs in Aging Glutamatergic Neurons

Glutamatergic neurons, responsible for excitatory signaling in the brain, play a key role in cognition, synaptic plasticity, and neural circuit function. We found that aging induces widespread chromatin and transcriptional changes in these neurons, with distinct regional vulnerabilities (Figure S6). In dentate gyrus (DG) neurons, age-related chromatin remodeling exhibits clear regional specificity, with the anterior hippocampus (AHC) and posterior hippocampus (PHC) showing divergent patterns of accessibility changes. Specifically, differential accessibility analysis identified 3,495 upregulated and 3,505 downregulated cCREs in aged DG neurons, with clusters of age-upregulated peaks enriched in the anterior hippocampus (AHC), while a distinct set of peaks showed greater downregulation in the posterior hippocampus (PHC) (Figure 4a). Motif enrichment analysis highlights regional differences in transcription factor activity, with CTCF motifs preferentially enriched in age-upregulated cCREs in the anterior hippocampus, suggesting altered chromatin looping and genome architecture in this region. Stress-responsive transcription factor complex AP-1 (Activator Protein 1) motifs are significantly upregulated in both brain regions (Figure 4b). Conversely, MEF2 motifs were strongly enriched in downregulated cCREs in the posterior hippocampus, suggesting a regional loss of transcriptional programs involved in synaptic plasticity and neuronal maintenance. MEF2 transcription factors regulate activity-dependent synapse remodeling and neuronal survival^40,42^.

**Figure 4.**
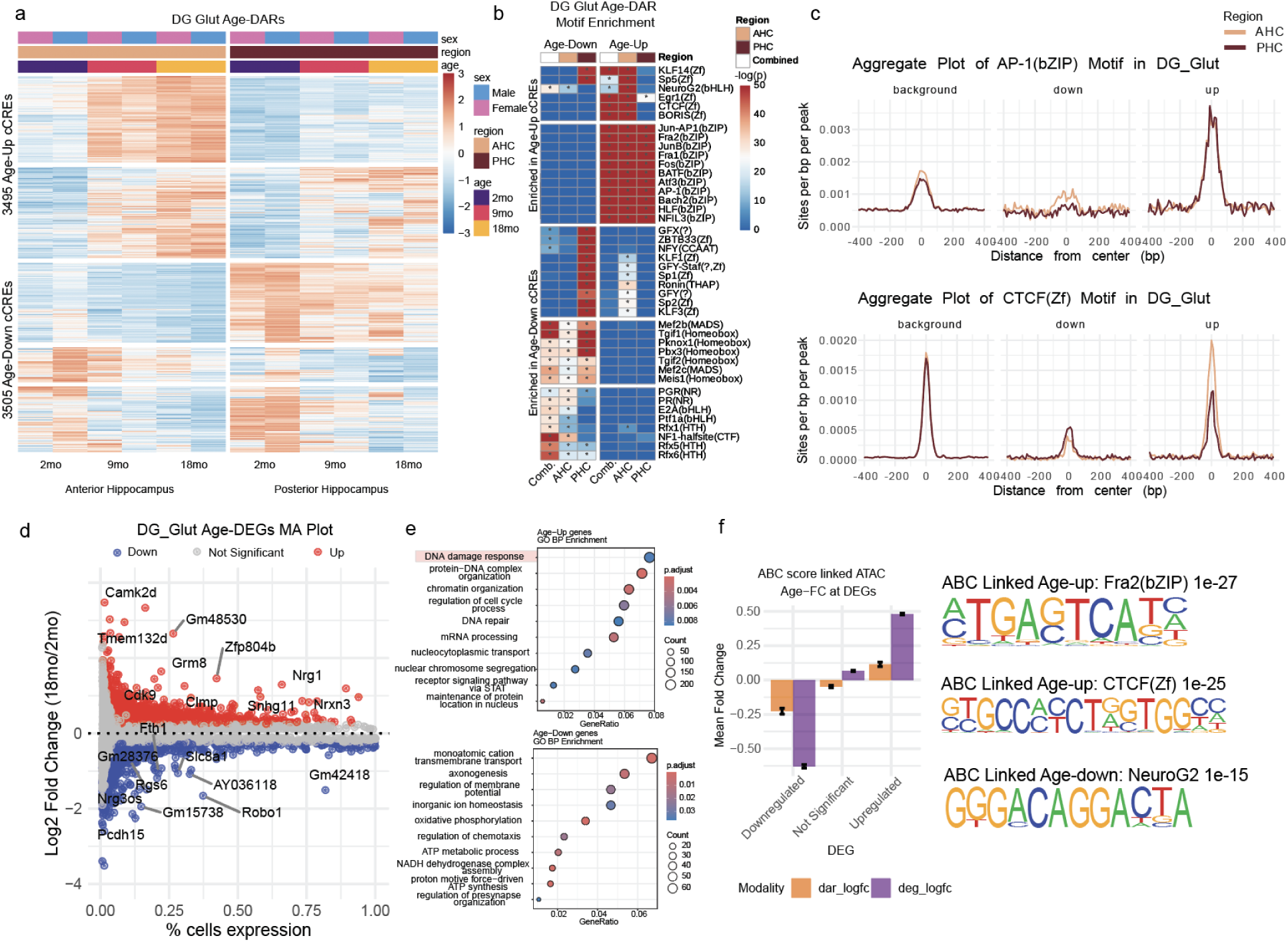
Aging induces region-specific regulatory reprogramming in dentate gyrus glutamatergic neurons. (A) Heatmap of normalized chromatin accessibility (z-score) at age-differential cCREs in dentate gyrus (DG) neurons across brain regions. (B) Motif enrichment analysis of downregulated (left) and upregulated (right) age-differential cCREs using all peaks as background; asterisks denote significance (adjusted p < 0.01). (C) Aggregate accessibility plots of AP-1 and CTCF motifs at upregulated, downregulated, and all cCREs across DG neuron regions. (D) MA plot of age-differential gene expression between 2-month-old and 18-month-old DG neurons. (E) Gene Ontology (GO) enrichment analysis of genes upregulated with age. (F) Bar plot showing mean age log_2_ fold change of downregulated, non-significant, and upregulated genes, along with corresponding changes in linked cCREs from ABC analysis. Right: Motif logos for the most enriched motifs in peaks linked to up- and downregulated genes.

Genes involved in DNA damage response are upregulated in aging DG neurons while genes involved in axonogenesis are downregulated (Figure 4d,e). ABC analysis reveals that open chromatin regions linked to age-upregulated genes in these neurons are enriched for AP1 and CTCF motifs, while those associated with downregulated genes are enriched for NeuroG2 motifs (Figure 4f). This indicates a shift from developmental transcriptional programs toward stress-responsive regulatory networks during aging, where AP1 activation is a key driver of aging-related gene expression changes. This is consistent with recent findings demonstrating that AP-1 drives chromatin opening at stress-responsive cis-regulatory elements while cell identity transcription factor binding sites decline^7^. These findings highlight the regional specificity of chromatin remodeling in DG neurons, where the anterior hippocampus exhibits stronger stress responses and chromatin reorganization, while the posterior hippocampus experiences a greater loss of neurodevelopmental regulatory programs.

Beyond the hippocampus, glutamatergic neurons across multiple brain regions undergo extensive chromatin and transcriptional changes with aging. Motif enrichment analysis highlights an increase in open chromatin elements recognized by stress-responsive transcription factors, particularly KLF and AP1, suggesting that aging is associated with enhanced stress adaptation in glutamatergic neurons (Figure S6b). Conversely, MEF2 and RFX motifs are enriched in downregulated cCREs across aging neurons, indicating a loss of transcription factor binding at sites critical for neuronal development and synaptic maintenance (Figure S6b). CTCF motifs show strong enrichment in age-upregulated peaks across multiple regions, including the frontal cortex (FC), entorhinal cortex (ENT), and anterior hippocampus (AHC), further supporting widespread chromatin remodeling during aging (Figure S6b). Gene Set Enrichment Analysis (GSEA) reveals consistent downregulation of axonogenesis and oxidative phosphorylation pathways, suggesting a decline in synaptic plasticity and metabolic function across multiple glutamatergic neuron subtypes (Figure S12b, S13b). These transcriptional shifts align with previous findings that neuronal aging is marked by reduced bioenergetic capacity and neurogenesis^3^. Together, these findings demonstrate that glutamatergic neurons undergo significant age-related chromatin reorganization and transcriptional remodeling, with region-specific regulatory shifts in DG neurons and widespread stress responses across cortical regions.

### Protocadherin Locus Derepression and Chromatin Remodeling in Aging Medium Spiny Neurons

GABAergic neuron populations play essential roles in inhibitory signaling across brain regions. With the exception of D12 medium spiny neurons (MSNs), which exhibit significant chromatin remodeling, most GABAergic subtypes show relatively modest chromatin and transcriptional changes with aging, though this may partially reflect lower cell numbers in our dataset (Figure S5a).

D12 medium spiny neurons (MSNs) are a key GABAergic subclass within the basal ganglia. This cell type is present in the caudate putamen (CP) and nucleus accumbens (NAC). We observed pronounced chromatin remodeling with aging in these cells. We further identified clusters of cCREs that are upregulated to higher levels in NAC and clusters of peaks more strongly downregulated in CP (Figure S9a). RFX motifs, which are involved in neuronal function^43^ and are broadly enriched in downregulated peaks of many neuronal subtypes, are instead enriched in upregulated peaks of NAC (Figure S9b). AP-1 motifs show significant enrichment in upregulated peaks across both regions, mirroring broader trends in glutamatergic neurons, where AP-1 activation is associated with stress responses and transcriptional dysregulation (Figures S9b, S6b). Notably, the AP-1 motif signal is strongest in D12 MSNs, suggesting these neurons experience particularly high levels of aging-related stress.

Differential gene expression analysis across age revealed genes related to cell junction organization were significantly enriched in upregulated genes in aging, including Neuregulin (Nrg) family members and protocadherins such as *Pcdhgc5* (Figure S9c). One of the most striking chromatin changes in aging D12 MSNs is increased accessibility at the Protocadherin gamma (Pcdhg) and Protocadherin alpha (Pcdha) loci, gene clusters essential for neuronal self-recognition and synaptic organization^44^. While protocadherins exhibit differential expression across multiple cell types, D12 MSNs show the strongest alterations at this locus, present in both brain regions (Figure S5f). Notably, transcriptional upregulation within the Pcdhg cluster is largely intronic, suggesting that regulatory elements or noncoding RNAs within this region may be differentially expressed rather than the protein-coding genes themselves. Among these, *3222401L13Rik*, a long noncoding RNA within the Pcdhg locus that has recently been implicated in aging astrocytes^45^, is upregulated across multiple cell types. 3C chromatin conformation data show structural changes at this locus, revealing increased contact frequency within the Pcdhg and Pcdha clusters while showing decreased interactions with upstream genomic regions (Figure S9e). These findings suggest that aging-induced chromatin reorganization in D12 MSNs may enhance protocadherin cluster activity while altering broader regulatory landscapes. The Pcdhg locus is part of a broader pattern of chromatin remodeling in aging neurons, where previously repressed genomic regions become more accessible. Recent studies have identified aging hotspots—genomic regions with significant accessibility gains—that overlap heterochromatin domains and are enriched in neurons^17^. In the next section, we examine the genome-wide consequences of heterochromatin loss, including transposable element activation, noncoding RNA upregulation, and stress-related reprogramming across cell types and brain regions.

### Heterochromatin Destabilization, Transposable Element Activation, and lncRNA Dysregulation in Aging Brains

Beyond cell-type-specific changes, we identified broader chromatin reorganization patterns associated with aging across multiple cell types. We next examined how heterochromatin domains and transposable elements are affected during brain aging. In our previous work, we found that aging is accompanied by widespread chromatin remodeling, particularly at heterochromatin domains and transposable elements (TEs)^17^, and aging neurons exhibit significant heterochromatin loss, particularly in cortical regions. With the increased resolution and broader scope of our current dataset spanning eight brain regions and multiple cell types, we now find that this phenomenon extends beyond neurons, with broad heterochromatin destabilization observed across diverse glial and neuronal populations. However, this effect remains most pronounced in glutamatergic neurons, particularly in the frontal cortex, hippocampus, entorhinal cortex, and amygdala, while the nucleus accumbens, periaqueductal gray, and caudate putamen exhibit little to no change (Figure S10).

To investigate heterochromatin alterations, we examined the overlap between age-associated differentially accessible regions and H3K9me3-marked heterochromatin domains from mouse forebrain^46^ (Methods). Age-upregulated peaks were significantly enriched in H3K9me3 domains, particularly in glutamatergic neurons (Figure 5a, middle), suggesting that aging is associated with the erosion of heterochromatin integrity. Consistent with this, transposable elements exhibited a pronounced shift toward increased accessibility across multiple cell types (Figure 5a, right), with specific subfamilies showing consistent patterns of activation across cell types (Figure S12c). This trend was evident in both male and female mice (Figure S10b), reinforcing heterochromatin erosion as a conserved feature of brain aging. Motif enrichment analysis revealed differentially upregulated regions within heterochromatin that were enriched for pioneer transcription factor AP1 family (bZIP)^47^ and CTCF motifs (Figure 5b).

**Figure 5.**
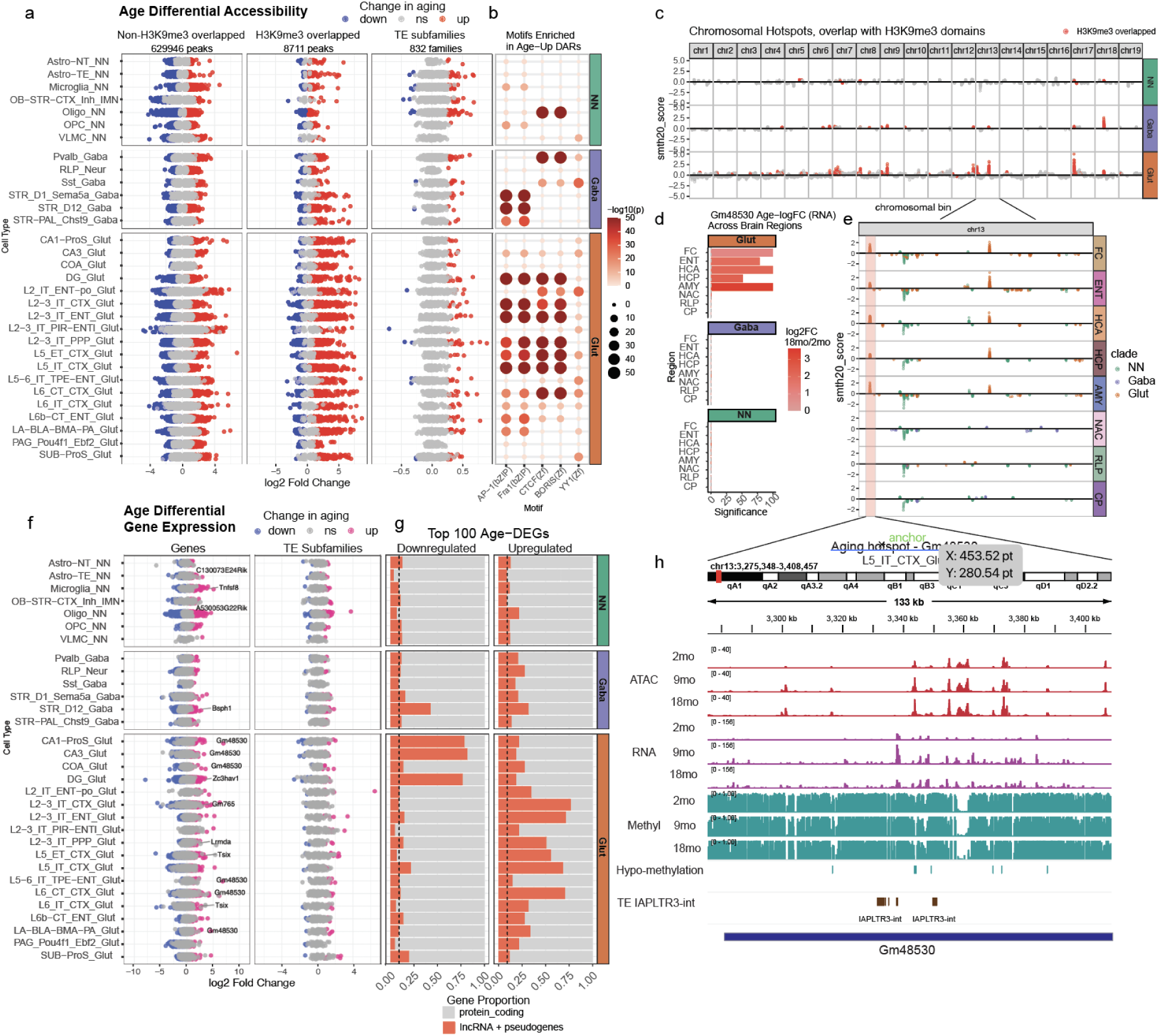
Upregulated accessibility and transcriptional shifts within heterochromatin-associated genomic hotspots. (A) Differentially accessible regions (DARs) in aging neurons. Left: Non-H3K9me3 peaks show a balanced distribution of up- and downregulated peaks. Middle: H3K9me3-overlapping peaks are predominantly upregulated, especially in glutamatergic neurons. Right: Transposable element (TE) subfamilies show widespread age-related increases in accessibility. (B) Motif enrichment analysis of age-upregulated DARs across neuronal populations. (C) Genomic hotspots with higher-than-expected density of DARs across cell types, colored by overlap with H3K9me3 domains. (D) log_2_ fold change and significance of lncRNA *Gm48530* expression across brain regions, located within an aging hotspot. (E) Chromosomal hotspot on chromosome 13 across brain regions; each point represents a cell type, colored by cell type clade. (F) Age-related transcriptional changes from scRNA-seq. Left: log_2_ fold change of genes across cell types, highlighting *Gm48530* among the most significantly upregulated. Right: TE subfamily expression shows a broader trend of upregulation. (G) Proportion of the top 100 most upregulated and downregulated genes that are long noncoding RNAs or pseudogenes. (H) Genome browser view of the chromosome 13 hotspot, including *Gm48530* and IAPLTR3-int, showing increased accessibility, expression, and hypomethylation with age.

Using Gaussian density scoring, a computational approach to identify genomic regions with significantly higher-than-expected concentrations of differential accessibility^17^ (Methods), we identified chromosomal aging hotspots across the genome. These hotspots were particularly upregulated in glutamatergic neurons, especially in cortical and hippocampal regions (Figure 5c). Notably, 47% of significantly upregulated hotspots overlap H3K9me3-marked domains, despite these regions covering only ~3% of the genome. This strong association between aging-associated accessibility gains and constitutive heterochromatin suggests a widespread breakdown of repressive chromatin compartments during aging. Many of these hotspots correspond to intergenic regions with TE activities, gene clusters that are typically repressed in mature neurons, and lncRNAs (Figure S3). Notably, lncRNAs were overrepresented among the most differentially expressed genes in aging, with over 50% of the top 100 upregulated genes in some cortical neuron cell types belonging to this category (Figure 5g). One of the most prominent aging-associated hotspots is located at the start of chromosome 13, where accessibility gains were observed across five brain regions (Figure 5e). This region contains the lncRNA *Gm48530*, which was the most significantly upregulated gene in aging cortical neurons across the same five brain regions (Figure 5d). The hotspot also overlaps *IAPLTR3-int*, a TE element that exhibited increased accessibility in multiple cell types (Figure 5h, S12c). Together, this locus exhibited coordinated increases in chromatin accessibility, gene expression, and hypomethylation (Figure 5h), suggesting that age-related chromatin remodeling at heterochromatin domains may alter regulatory landscapes and lead to transcriptional activation.

Beyond autosomal hotspots, several aging-associated hotspots were found in the X chromosome in female samples (Figures S11a, S3). The most significant of these occurred at the Firre lncRNA locus, which exhibited continuous and drastic accessibility increases (Figure S11a). This finding aligns with previous reports linking aging to the loss of X chromosome inactivation^48–50^. Similarly, the lncRNA *Dxz4*, another key element involved in X chromosome organization, exhibited increased accessibility in aging neurons, suggesting that large-scale shifts in X-linked chromatin architecture may accompany aging (Figure S11a). In parallel, Xist and Tsix, two critical regulators of X chromosome inactivation, were among the most differentially-upregulated genes across multiple female brain regions and cell types, especially in neurons, consistent with previous research (Fig. S11c)^49^. This could be a compensatory mechanism to counteract the possible loss of X chromosome inactivation we observe. Interestingly, while the *Firre* and *Dxz4* loci gained accessibility with age, the pseudoautosomal region at the end of the X chromosome, which typically escapes X inactivation, exhibited a striking loss of chromatin accessibility and gene expression in female mice (Figure S11a, b). The most affected region included *Erdr1* (*Gm47283*), which was the most consistently downregulated lncRNA in female brain aging across cell types and brain regions. This opposing pattern—decreased accessibility at normally active X-linked regions alongside increased accessibility at heterochromatin-associated loci - suggests that chromosome X undergoes extensive chromatin reorganization in aging females, potentially reflecting an extension of global heterochromatin destabilization.

Currently, the mechanisms responsible for the aging associated heterochromatin decay remain unclear. We noted that among the most consistently down-regulated pathways across cell types was oxidative phosphorylation (Figure S13a,b), highlighting the central role of metabolic decline in brain aging. This was particularly pronounced in neuronal populations, where energy demands are highest^51^. Genes involved in protein synthesis were also broadly downregulated, consistent with reduced energy availability in these cells. Additionally, we observed downregulation of genes involved in heterochromatin maintenance (Figure S13c), including histone methyltransferases and chromatin remodeling factors, providing a potential mechanistic explanation for the heterochromatin instability observed across cell types. These findings highlight heterochromatin remodeling, TE activation, and lncRNA dysregulation as key hallmarks of aging in the brain. The widespread loss of heterochromatin integrity, coupled with transcriptional shifts in chromatin organization and increased TE activity, suggests that aging involves a progressive breakdown of epigenetic regulatory barriers. This dysregulation may contribute to genomic instability and functional decline in aging neurons, underscoring the need for further investigation into chromatin-based mechanisms of neurodegeneration.

## Discussion

We show here that aging is accompanied by profound changes in chromatin accessibility, transcription factor activity, and cell identity maintenance in the brain. Our multi-omic analysis across eight brain regions reveals key regulatory features underlying these changes, linking master transcription factor loss, stress-induced reprogramming, and heterochromatin instability to cellular dysfunction in aging.

A major hallmark of aging observed in our data is the decline of transcription factors crucial for cell identity and function. In progenitor populations, factors such as *Lhx2* and *Pax6* show significant reductions, impairing neurogenic capacity. Similarly, oligodendrocytes and their precursors (IOLs) exhibit downregulation of *Sox* genes, disrupting myelination and oligodendrocyte maintenance. In neurons, master transcription factors including *Mef2* and *NeuroG2* lose function, likely contributing to deficits in axonogenesis and synaptic plasticity, as seen in downregulated genes. As developmental transcription factors decline, aging cells exhibit an upregulation of stress-responsive transcription factors. We observe increased accessibility of motifs for AP1 (bZIP) in neurons, KLF across cell types, and IRF8 in microglia, linking aging to inflammation, oxidative stress, and immune activation.

Another key feature of aging shown in our data is the widespread loss of heterochromatin integrity, accompanied by dysregulation of transposable elements (TEs) and non-coding RNAs. Long noncoding RNAs (lncRNAs) are among the most upregulated genes, particularly those located within heterochromatin-associated domains. Aging hotspots overlapping heterochromatin regions are consistently observed across autosomes in both male and female mice. However, female mice uniquely exhibit increased chromatin accessibility on the X chromosome, alongside a concurrent loss of accessibility and expression at the pseudo-autosomal region. These findings suggest that aging-driven chromatin reorganization extends beyond autosomal heterochromatin erosion, affecting large-scale X-linked regulatory landscapes in a sex-specific manner. This dimorphism in X chromosome regulation may contribute to known sex differences in brain aging trajectories and neurodegeneration susceptibility, though the functional consequences require further investigation.

Our data suggest that aging is associated with a loss of transcriptional and epigenetic control over cell identity, causing cells to drift away from their defined states. The concurrent loss of developmental transcription factors, increased stress response activity, and heterochromatin breakdown contribute to a landscape where regulatory control is progressively lost. In this context, our findings align with emerging models of aging that propose a loss of epigenetic constraints leading to transcriptional drift^7,52^. The reactivation of TEs and lncRNAs, coupled with downregulation of genes important to metabolic and cellular function, suggests that aging cells fail to maintain repression of elements that should remain silenced while also struggling to sustain essential transcriptional programs critical for homeostasis and function.

Epigenetic and transcriptional interventions have shown promise in mitigating aging-related decline, and our findings provide a mechanistic context for these approaches. The widespread loss of heterochromatin integrity in aging neurons aligns with studies showing that reinforcing heterochromatin maintenance can extend lifespan and preserve cellular function^53,54^. Similarly, the depletion of lineage-defining transcription factors suggests that stabilizing these networks could counteract the progressive loss of cell identity with age. Reprogramming with pluripotency factors has been shown to reverse aspects of cellular aging^8,55^ yet the underlying mechanisms remain unclear. Our findings suggest that aging brain cells lose regulatory control, shifting from developmental transcription factor networks toward stress-related reprogramming. Future efforts should focus on targeted strategies to reinforce lost transcriptional programs and maintain heterochromatin, with the goal of preserving cognitive function and delaying neurodegeneration.

## Limitations of the study

This study captures key chromatin and transcriptional changes across aging but is limited to three timepoints, which may not fully resolve the temporal dynamics of regulatory decline. The 18-month timepoint reflects mid-to-late adulthood in mice, not extreme old age. Sex-specific analyses are constrained by differing methodologies between sexes (snATAC-seq in males, 10X Multiome in females), limiting direct comparisons. Finally, the findings are correlative; causal relationships between transcription factor dysregulation, chromatin remodeling, and aging phenotypes remain to be tested through perturbation studies.

## Resource Availability

### Lead Contact

Further information and requests for resources and reagents should be directed to Bing Ren and M Marga Behrens.

### Materials Availability

This study did not generate new unique reagents.

### Data and Code Availability

The raw data from this study have been deposited in SRA under project number PRJNA1248586. The processed data from this study have been deposited in the NCBI’s Gene Expression Omnibus (GEO) database under accession number GSE294772. Analysis code are available on GitHub at https://github.com/luisajamaral/aging_mouse_brain_code. Any additional information needed to reanalyze the data reported in this paper can be obtained from the lead contact upon request.

## Acknowledgments

This work was supported by the National Institutes of Health (NIH) grant 5R01AG066018 to B.R and J.R.E.

## Author Contributions

B.R., J.R.E., M.M.B., and M.L.A. conceived and designed the study. M.L.A. performed analysis and drafted the manuscript. J.Osteen, A.B., J.R., N.D.J., and S.Ca. contributed to tissue dissection and nuclei production. M.M., Q.Y., E.E., and E.W.S. generated the 10x Multiome data. X.H. generated the snATAC-seq data. A.W. oversaw data generation for scATAC and 10X Multiome data. C.T.B.B., J.A., S.C., N.E., and J.L. generated MERFISH data. J.Olness. and C.K. preprocessed the MERFISH data under the supervision of Q.Zhu. Q.Z. analyzed shared snm3c-seq data and calculated ABC scores. S.M. performed RNA doublet removal. S.Z. provided datasets for label transfer. M.L.A., S.M., and T.L. managed public data release. J.R.E., M.M.B., M.L.A., and B.R. coordinated the research and edited the manuscript. B.R. supervised the data collection and analysis.

## Supplementary Figures

**Figure S1.**
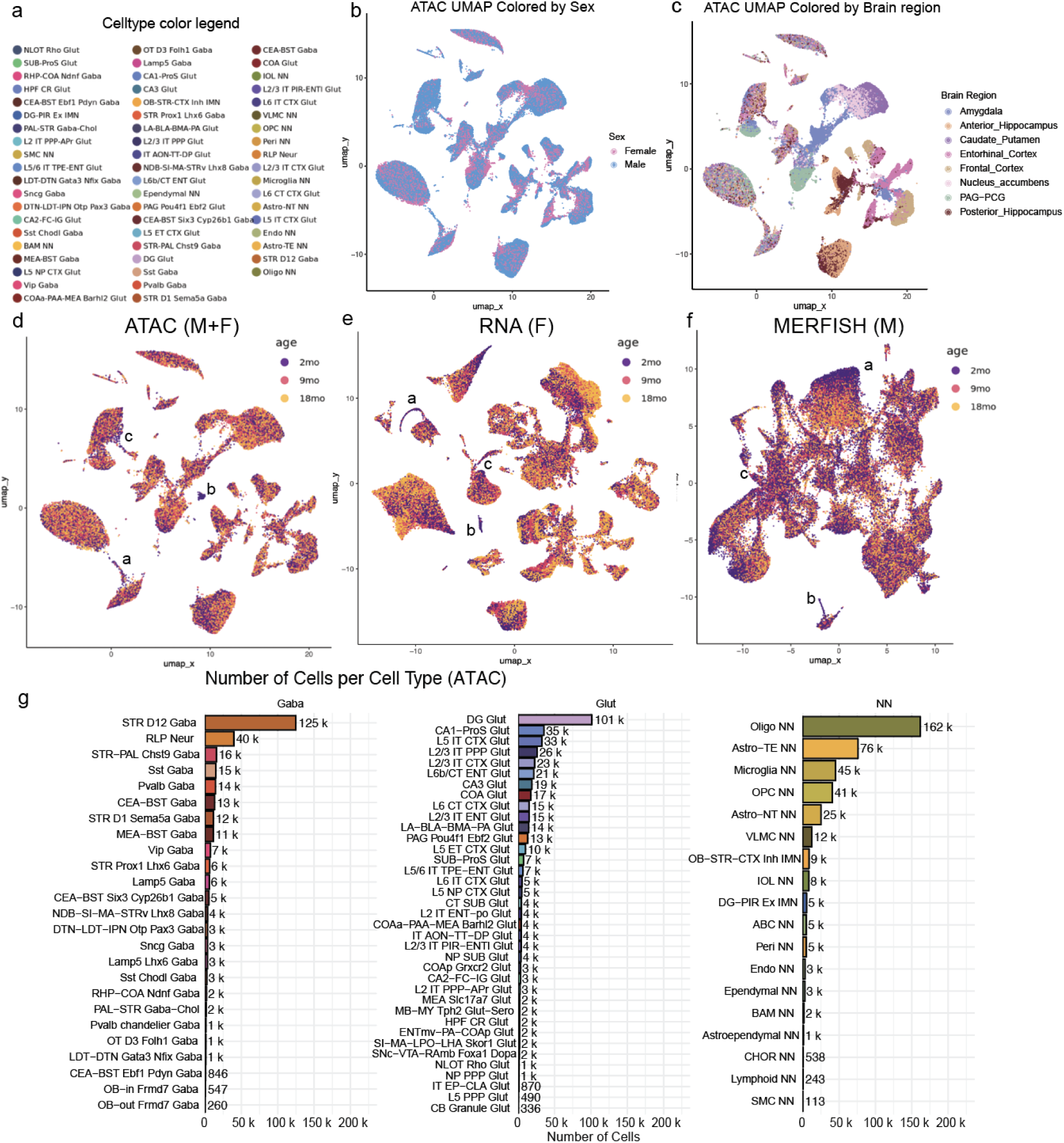
Characterization of multi-omic datasets and age-related gene regulation. (A) Cell type color legend corresponding to Figures 1B and 2A. (B)UMAP of snATAC-seq data annotated by sex. (C) UMAP of snATAC-seq data annotated by brain region. (D-F) UMAP projections of scATAC-seq, scRNA-seq, and MERFISH data colored by age group (2mo, 9mo, 18mo). (G) Number of cells per celltype for ATAC data (male and female)

**Figure S2.**
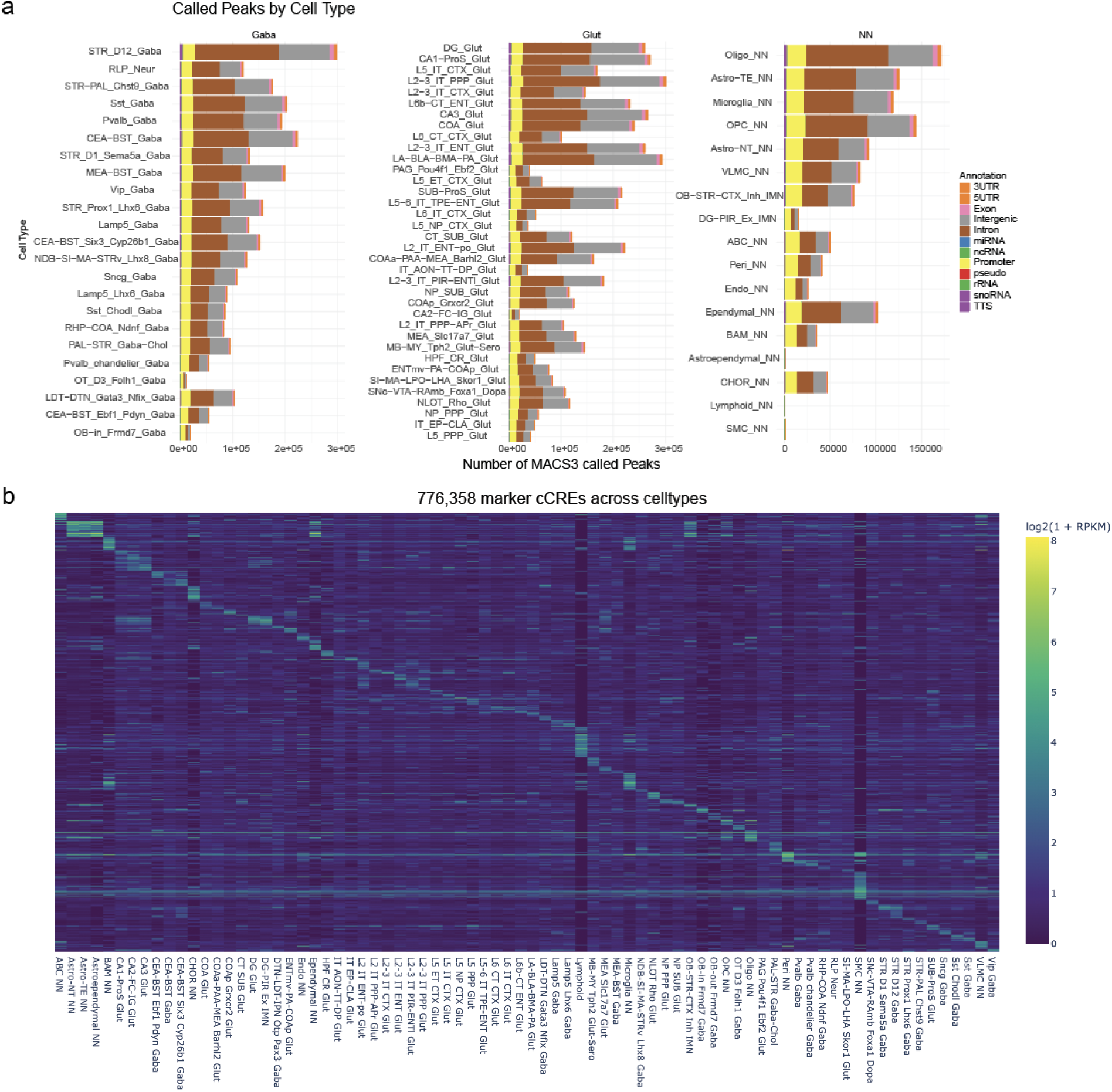
Cell type-level peak calling identifies cell-type-specific regulatory signatures. (A) Number of peaks called by MACS3 per cell type, colored by genomic annotation. (B) Heatmap showing accessibility (log_2_[1 + RPKM]) at marker cis-regulatory elements (cCREs) across cell types.

**Figure S3.**
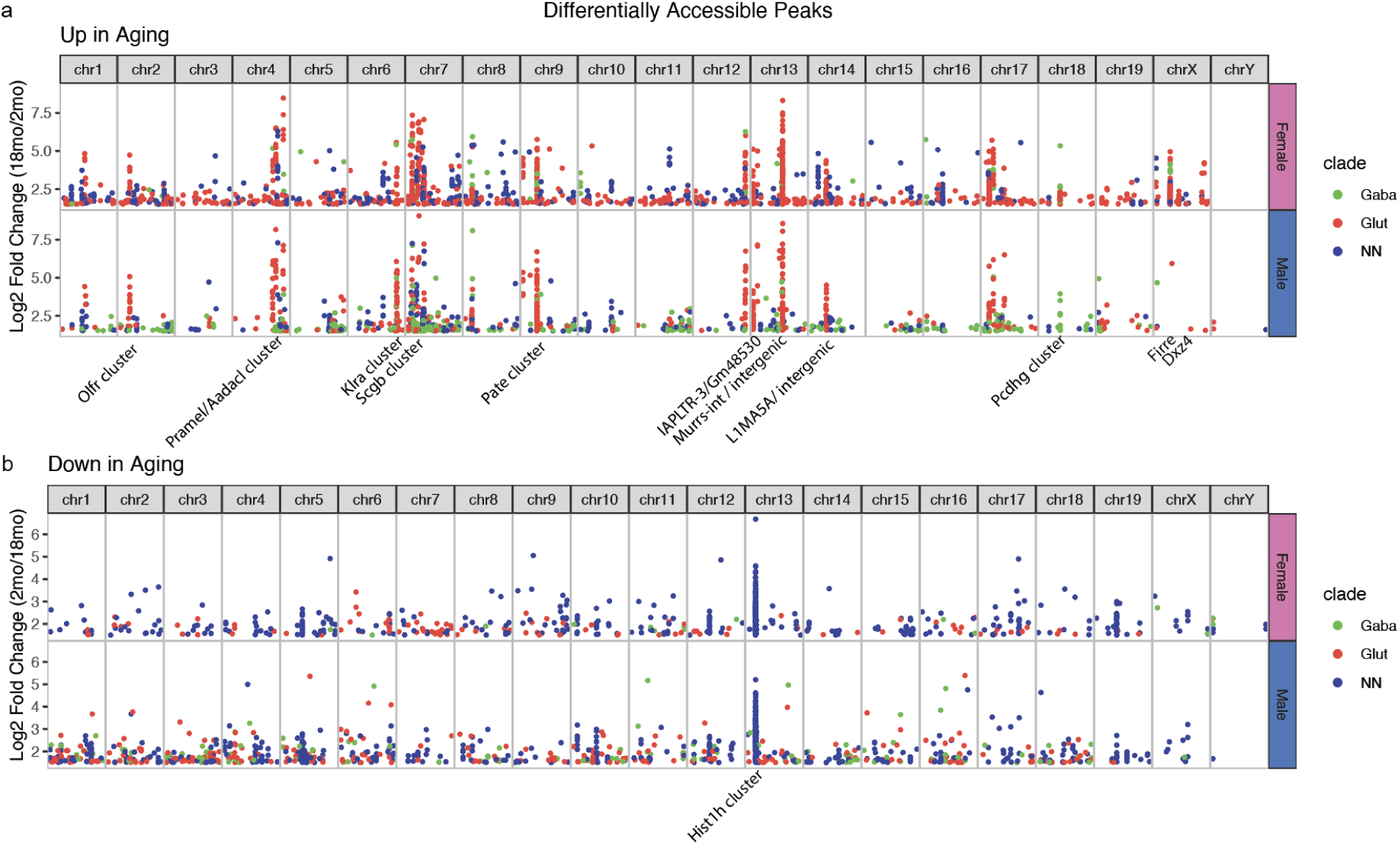
Genome-wide locations of age-associated chromatin accessibility changes in male and female brains. (A) Genomic locations of age-upregulated differentially accessible regions (DARs), plotted as log_2_ fold change (18mo/2mo) across chromosomes. (B) Genomic locations of age-downregulated DARs, plotted as log_2_ fold change (2mo/18mo) across chromosomes.

**Figure S4.**
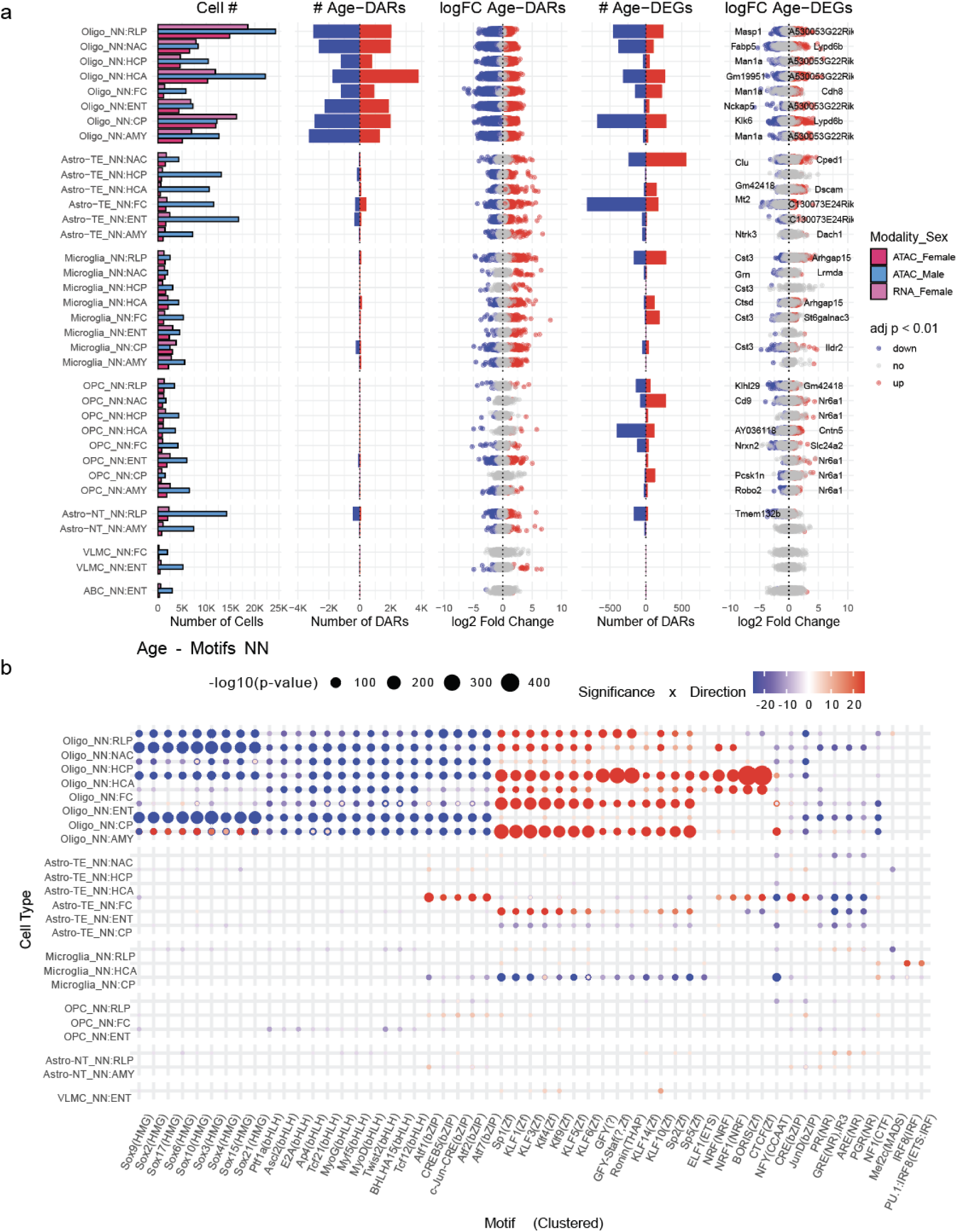
Aging-associated chromatin and transcriptional changes in non-neuronal cell types. (A) Left to right: Cell counts per non-neuronal cell type, brain region, and modality. Bar plot showing the number of age-associated DARs (adj. p < 0.01, |log_2_FC| > 0.25). Scatter plots of log_2_ fold change (log_2_FC) for DARs, colored by direction (up- or downregulated with age). Bar plot showing the number of age-associated DEGs (adj. p < 0.01, |log_2_FC| > 0.25). (B) Motif enrichment in age-DARs across non-neuronal cell types. Dot size indicates −log_10_(p-value), and color denotes direction of change with aging.

**Figure S5.**
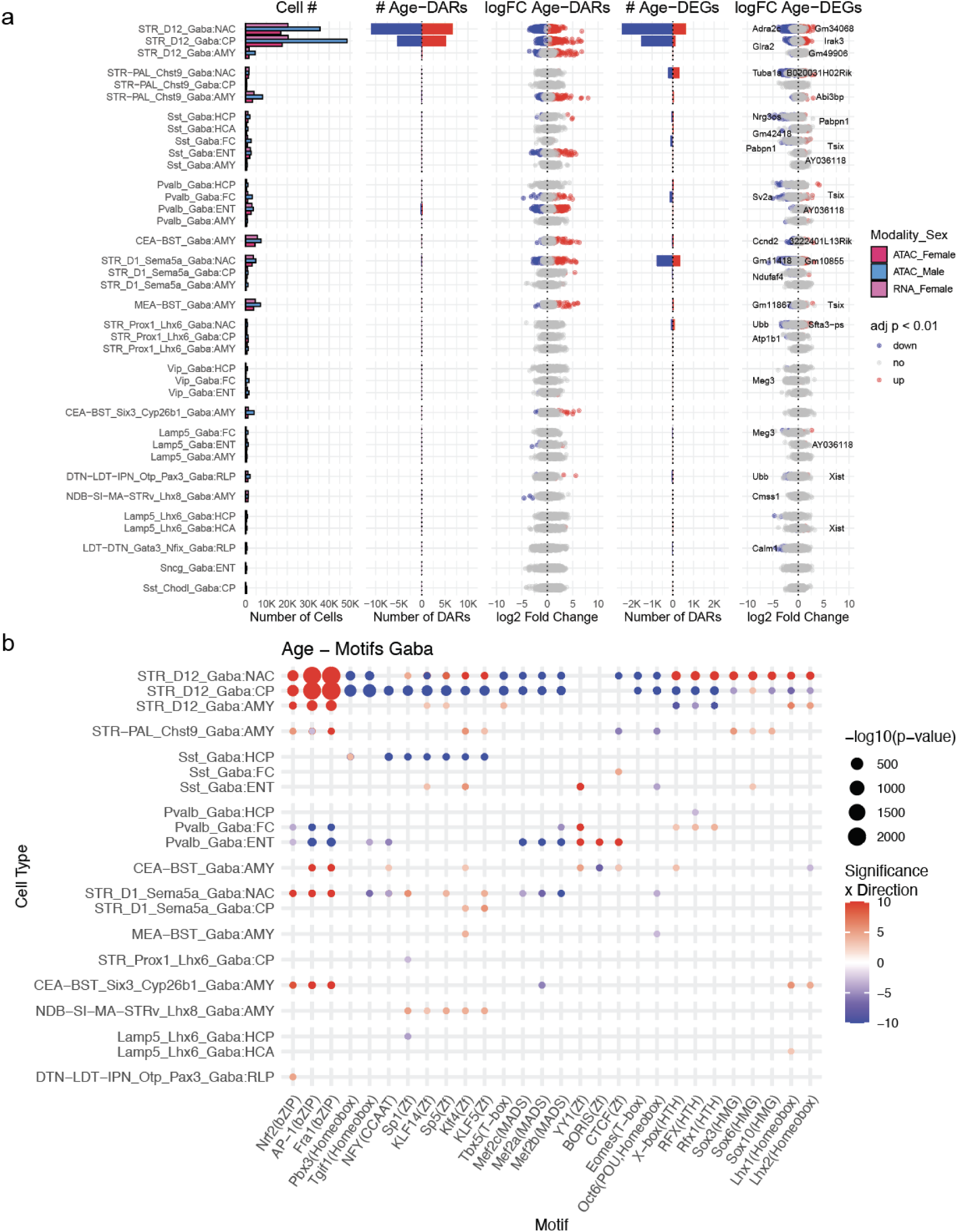
Aging-associated chromatin and transcriptional changes in GABAergic cell types. (A) Left to right: Cell counts per GABAergic cell type, brain region, and modality. Bar plot showing the number of age-associated DARs (adj. p < 0.01, |log_2_FC| > 0.25). Scatter plots of log_2_ fold change (log_2_FC) for DARs, colored by direction (up- or downregulated with age). Bar plot showing the number of age-associated DEGs (adj. p < 0.01, |log_2_FC| > 0.25). Scatter plots of log_2_FC for DEGs, colored by direction of change. (B) Motif enrichment in age-DARs across GABAergic cell types. Dot size indicates −log_10_(p-value), and color denotes direction of change with aging.

**Figure S6.**
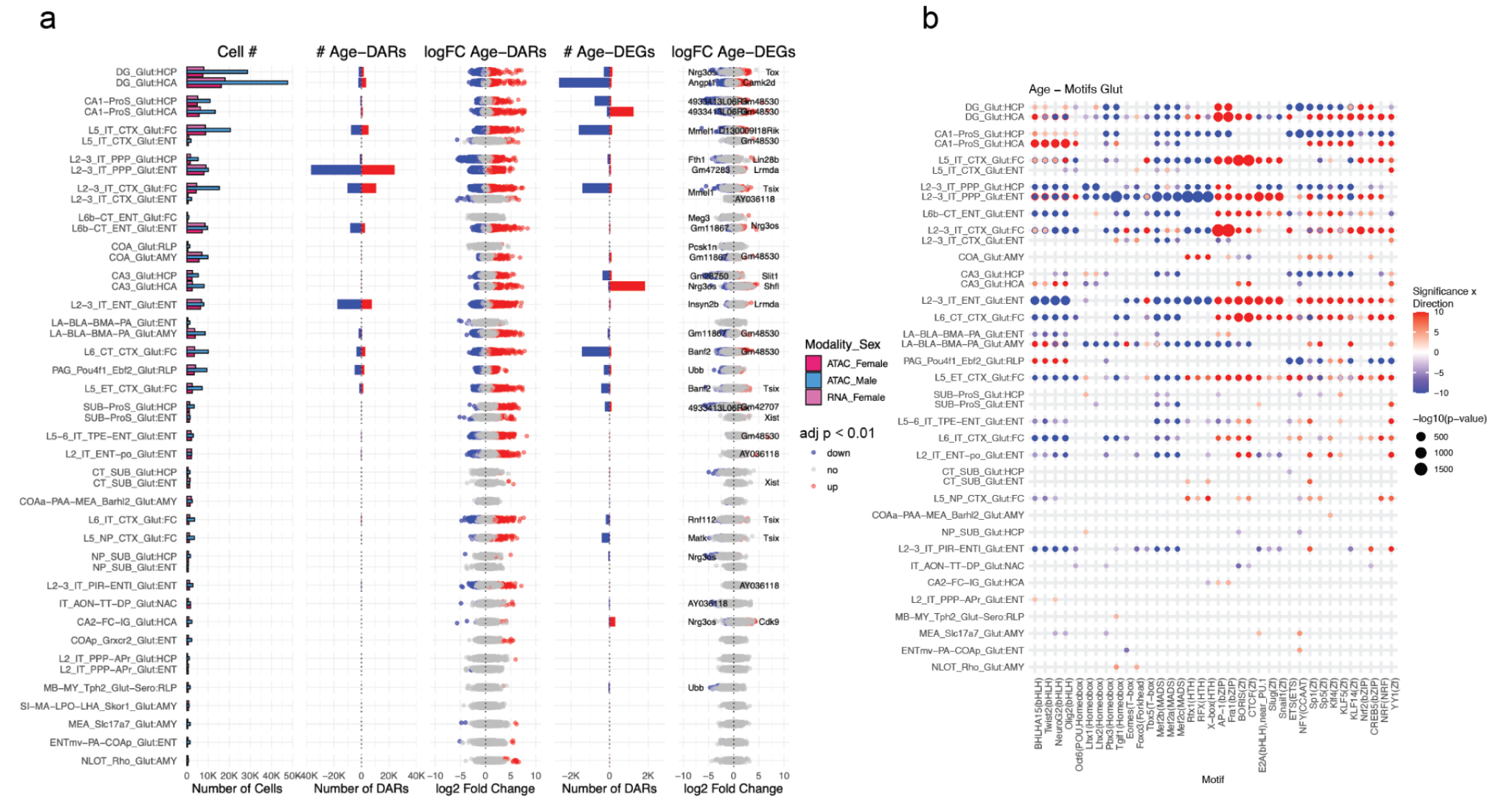
Aging-associated chromatin and transcriptional changes in glutamatergic cell types. (A) Left to right: Cell counts per glutamatergic cell type, brain region, and modality. Bar plot showing the number of age-associated DARs (adj. p < 0.01, |log_2_FC| > 0.25). Scatter plots of log_2_ fold change (log_2_FC) for DARs, colored by direction (up- or downregulated with age). Bar plot showing the number of age-associated DEGs (adj. p < 0.01, |log_2_FC| > 0.25). Scatter plots of log_2_FC for DEGs, colored by direction of change. (B) Motif enrichment in age-DARs across glutamatergic cell types. Dot size indicates −log_10_(p-value), and color denotes direction of change with aging.

**Figure S7.**
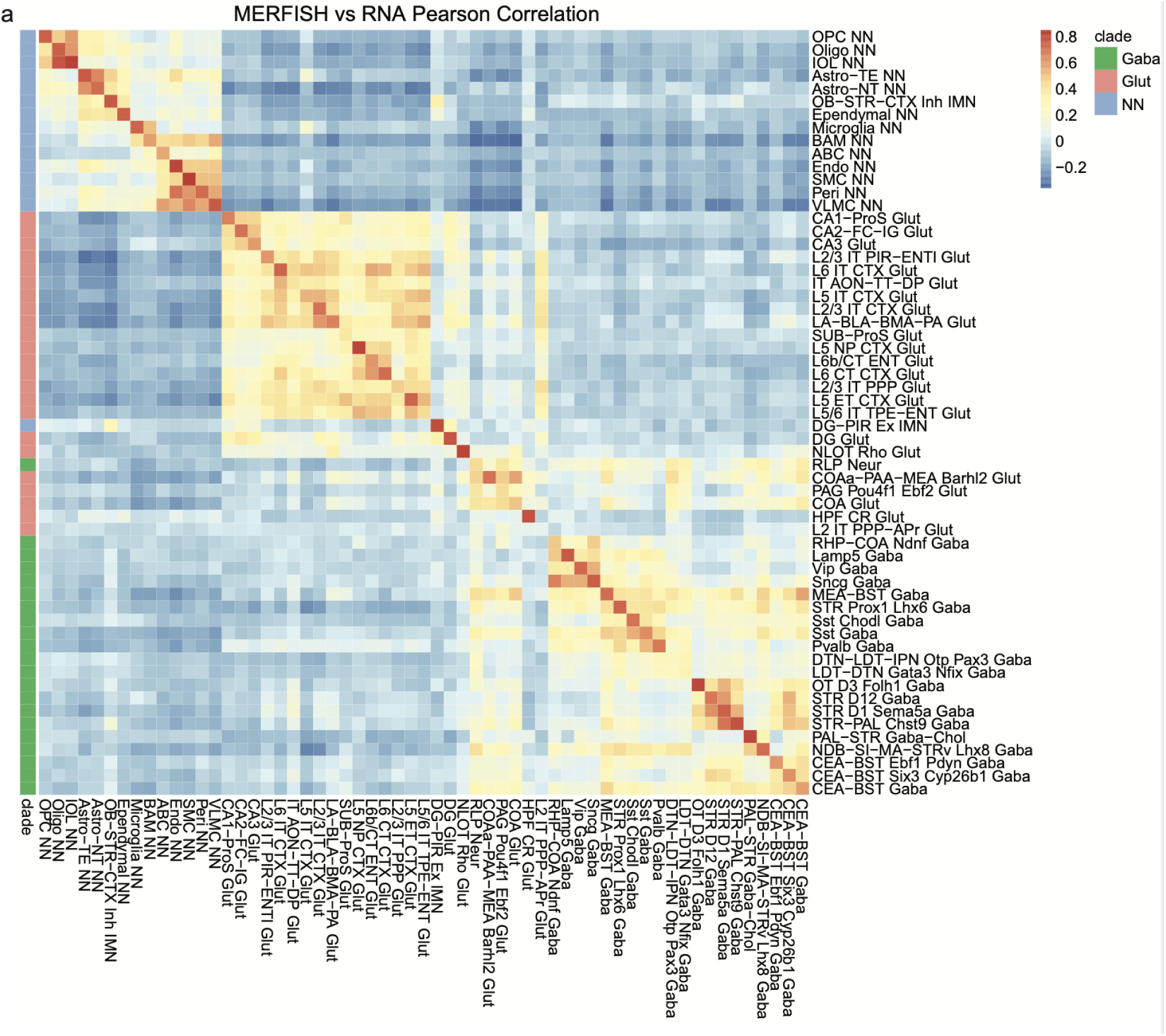
Cross-modal correlation of gene expression across cell types. (A) Heatmap showing Pearson correlation coefficients between RNA-seq and MERFISH gene expression profiles across shared cell types. Expression values were CLR-normalized per cell, aggregated by cell type, and RNA-seq values were additionally normalized by gene length. Correlations were computed across shared genes to assess concordance between modalities.

**Figure S8.**
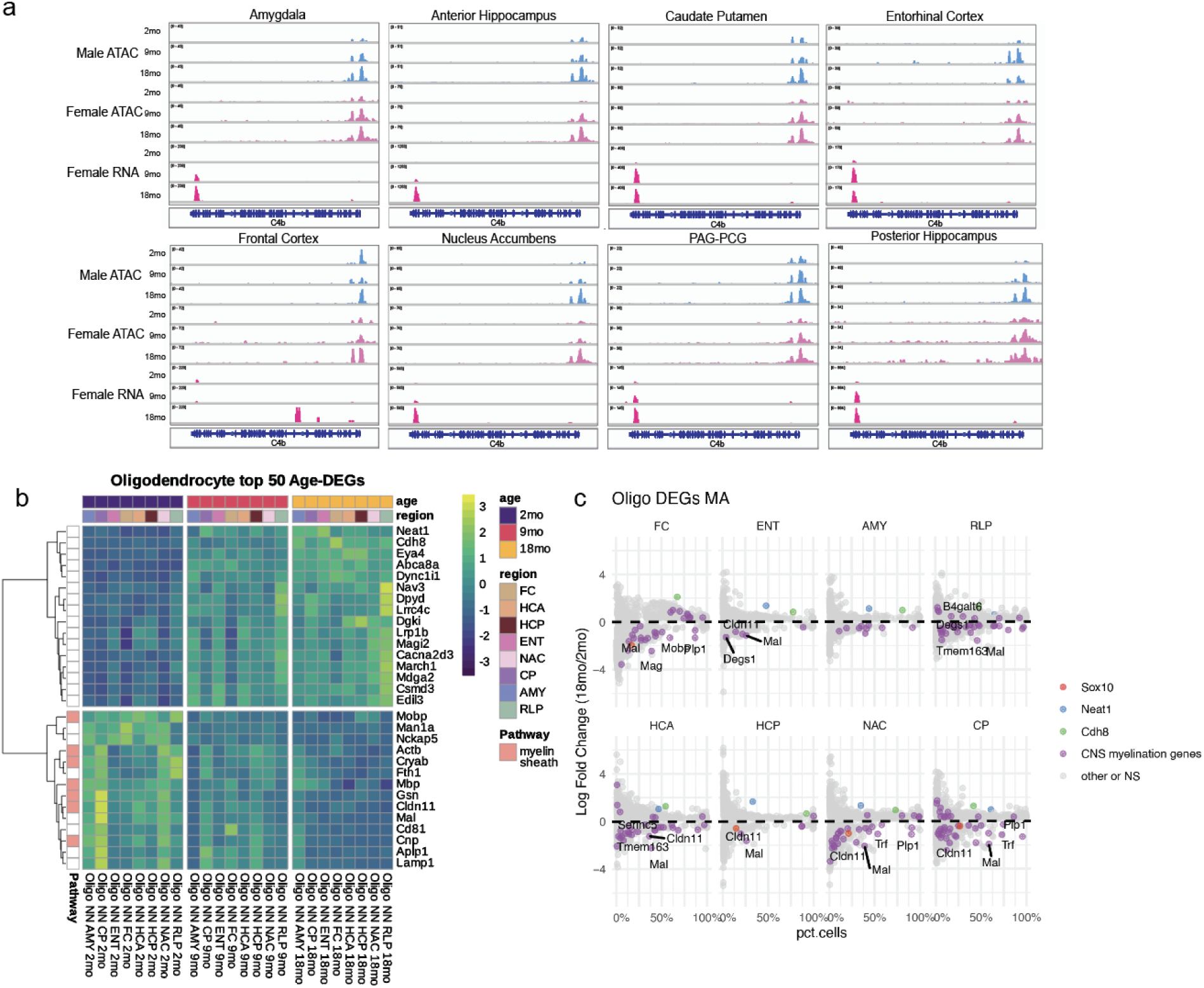
Downregulation of myelination-associated genes in aging oligodendrocytes. (A) Genome browser tracks showing chromatin accessibility (male ATAC, female ATAC) and RNA expression (female RNA) across age in eight brain regions in oligodendrocytes at the age-associated gene C4b. (B) Heatmap of the top 30 age-associated DEGs in oligodendrocytes, showing consistent downregulation of genes enriched for the “myelin sheath” GO term across brain regions. (C) MA plots of DEGs in oligodendrocytes, highlighting myelination-related genes (*Mal*, *Mog*, *Olig1*). X-axis denotes the percentage of cells expressing each gene; Y-axis shows log_2_ fold change (18mo vs. 2mo).

**Figure S9.**
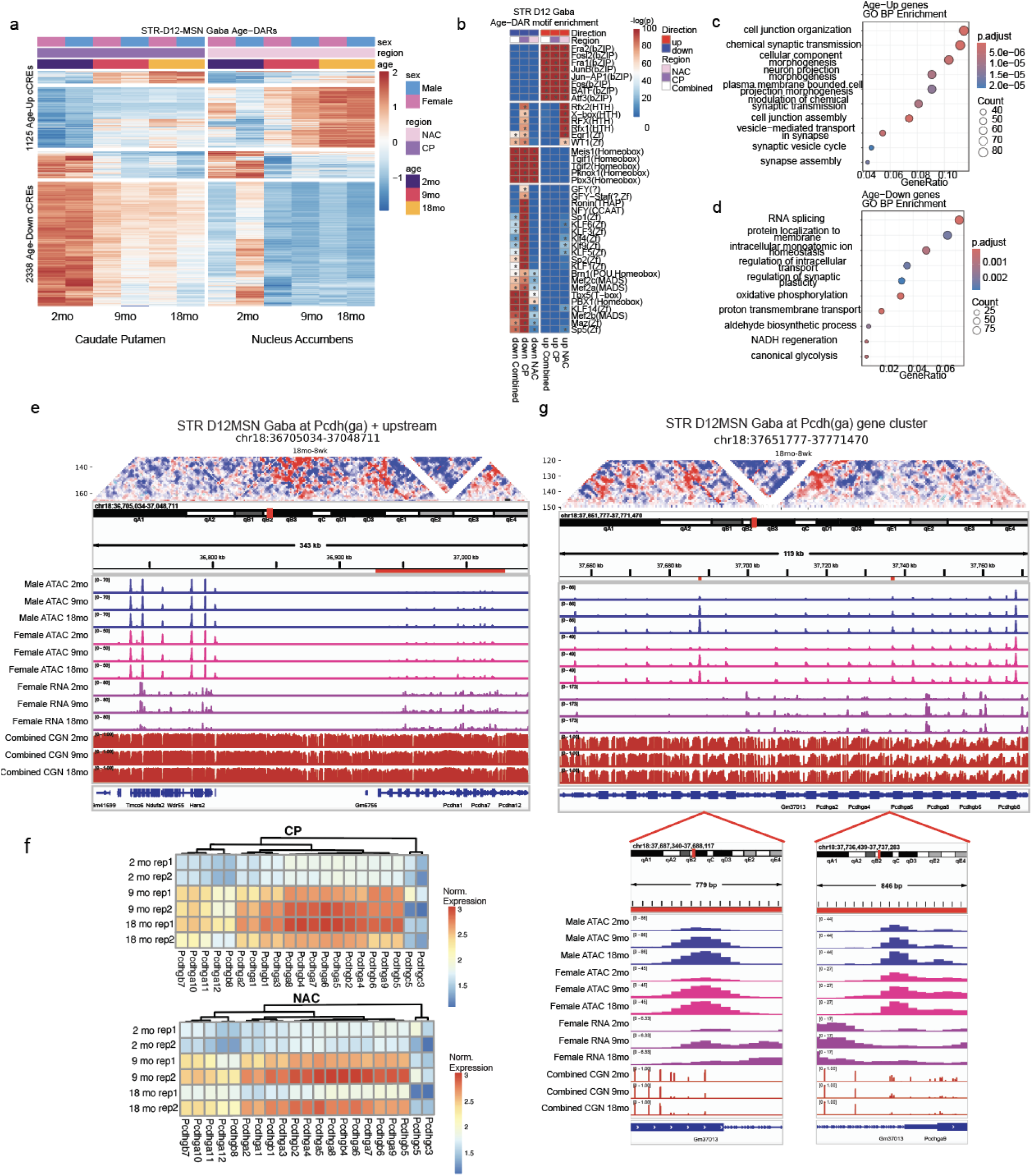
Aging-associated chromatin remodeling and protocadherin locus reorganization in D12 MSN neurons. (A) Heatmap showing scaled, normalized chromatin accessibility at age-differential cCREs in D12 medium spiny neurons (MSNs) from the striatum. (B) Motif enrichment analysis for age-upregulated and age-downregulated cCREs in D12 MSNs. (C) MA plot of age-differentially expressed genes (DEGs) in D12 MSNs (18mo vs. 2mo). (D) GO Biological Process enrichment for age-upregulated (left) and age-downregulated (right) DEGs. (E) Genome browser view of the *Protocadherin gamma* (Pcdhg) gene cluster in D12 MSNs showing increased chromatin accessibility, gene expression, hypomethylation, and altered Hi-C contact frequency with aging. (F) Heatmap of normalized expression levels for *Pcdhg* genes across age groups and brain regions in D12 MSN GABAergic neurons. (G) Zoomed-in IGV view of the *Pcdhg* locus highlighting specific aging-associated regulatory changes.

**Figure S10.**
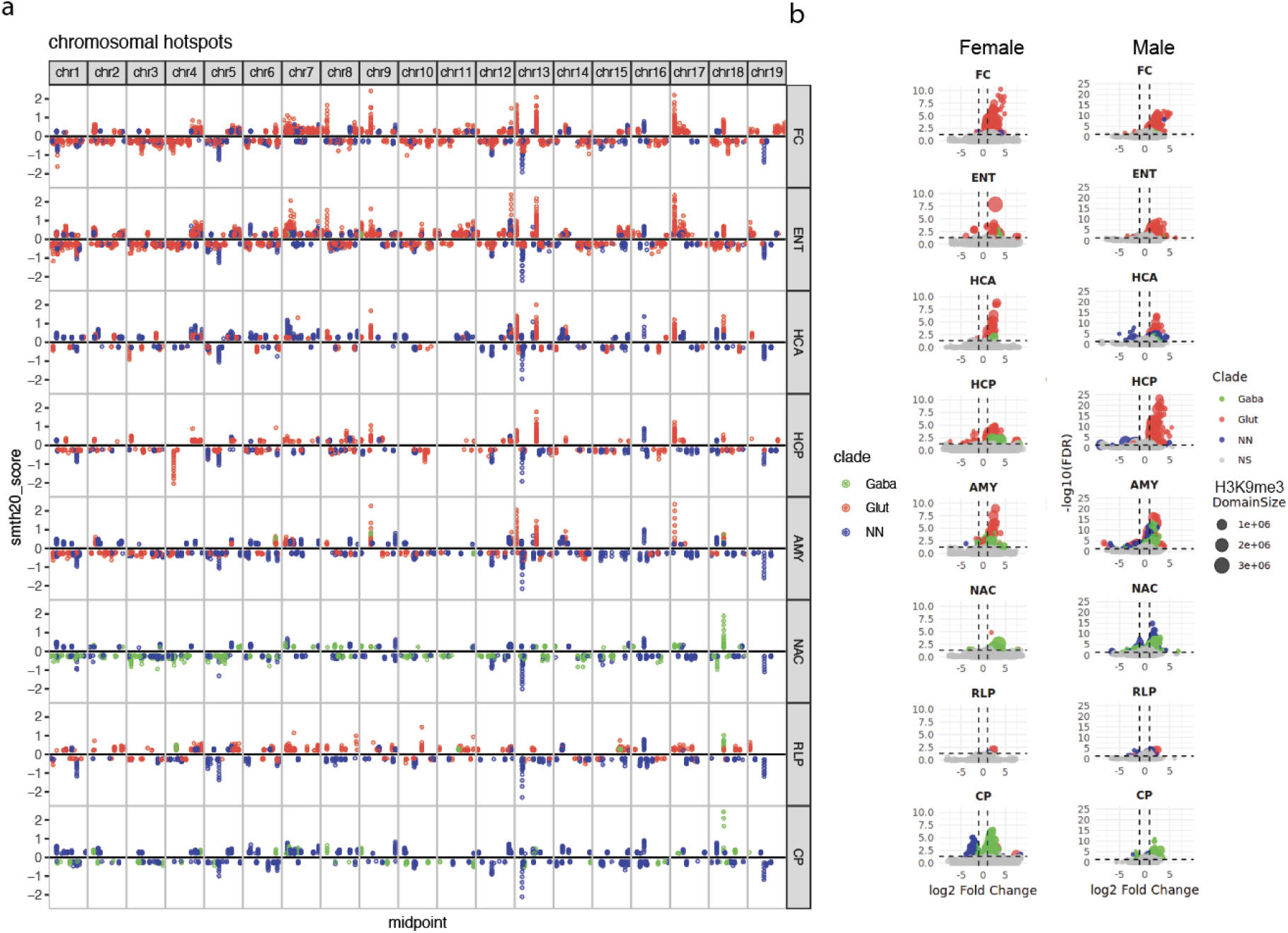
Chromosomal aging hotspots and heterochromatin loss across brain regions. (A) Chromosomal aging hotspots with significantly more DARs than expected by chance across cell types. Points are colored by cell type clade (glutamatergic, GABAergic, non-neuronal) and faceted by brain region, highlighting regions of age-associated chromatin accessibility gain or loss. (B) Volcano plots of differential chromatin accessibility within H3K9me3-marked domains across brain regions for glutamatergic, GABAergic, and non-neuronal cells. X-axis: log_2_ fold change (18mo/2mo); Y-axis: −log_10_(adjusted p-value). Point size represents the length of the H3K9me3 domain.

**Figure S11.**
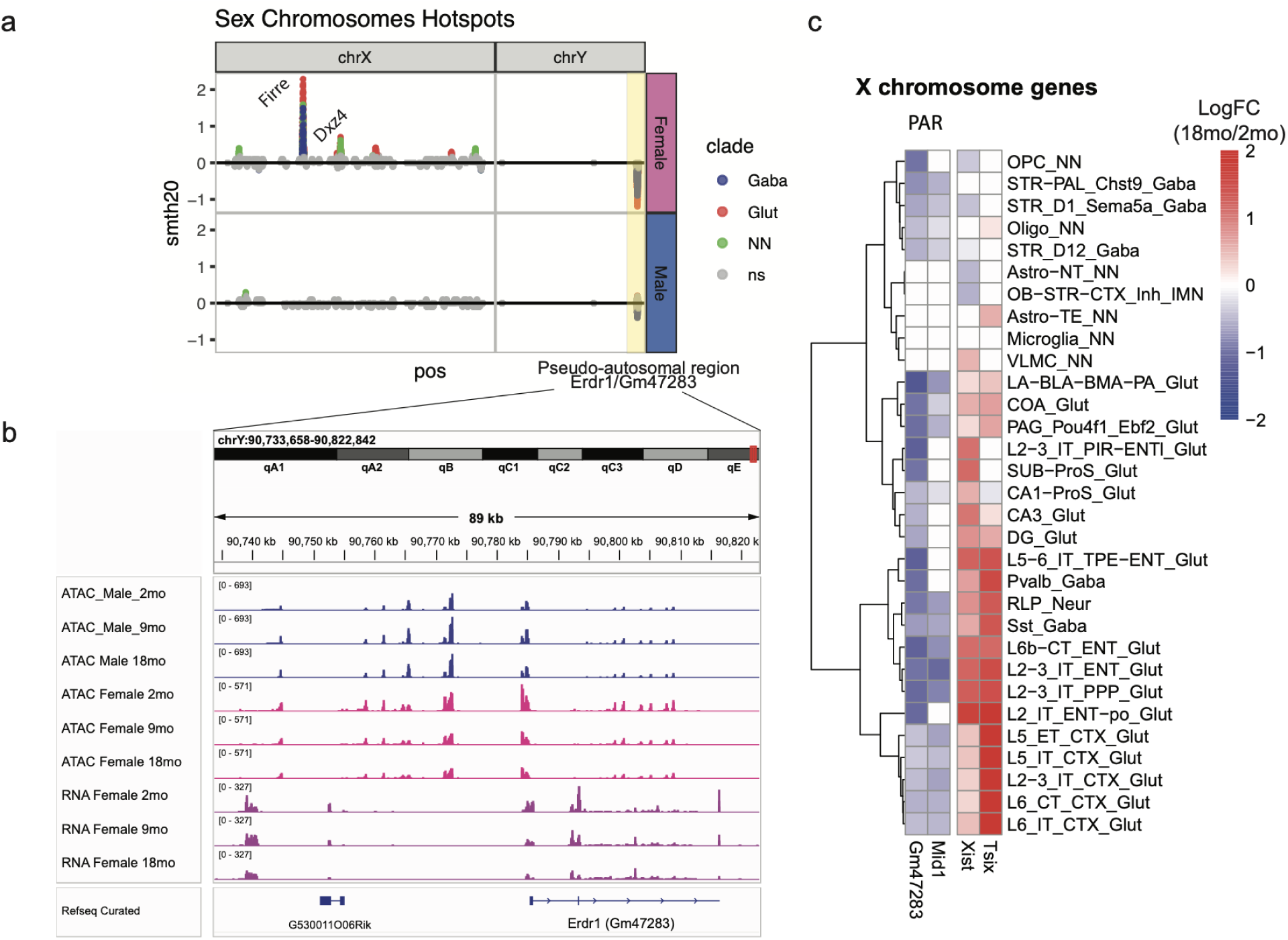
Sex chromosome aging hotspots and pseudoautosomal region remodeling. (A) Aging-associated chromatin accessibility hotspots on the sex chromosomes. Highlighted region at the distal end of chromosome Y corresponds to the pseudoautosomal region. (B) Genome browser view (IGV) of the pseudoautosomal region containing *Erdr1* (also known as *Gm47283*) across representative cell type STR_D12_GABA, showing chromatin accessibility (male ATAC, female ATAC) and RNA expression (female RNA) across age groups. (C) Heatmap of age-related log_2_ fold change (18mo/2mo) in expression for pseudoautosomal region genes (*Gm47283*, *Mid1*) and X-inactivation regulators (*Xist*, *Tsix*) across major cell types.

**Figure S12.**
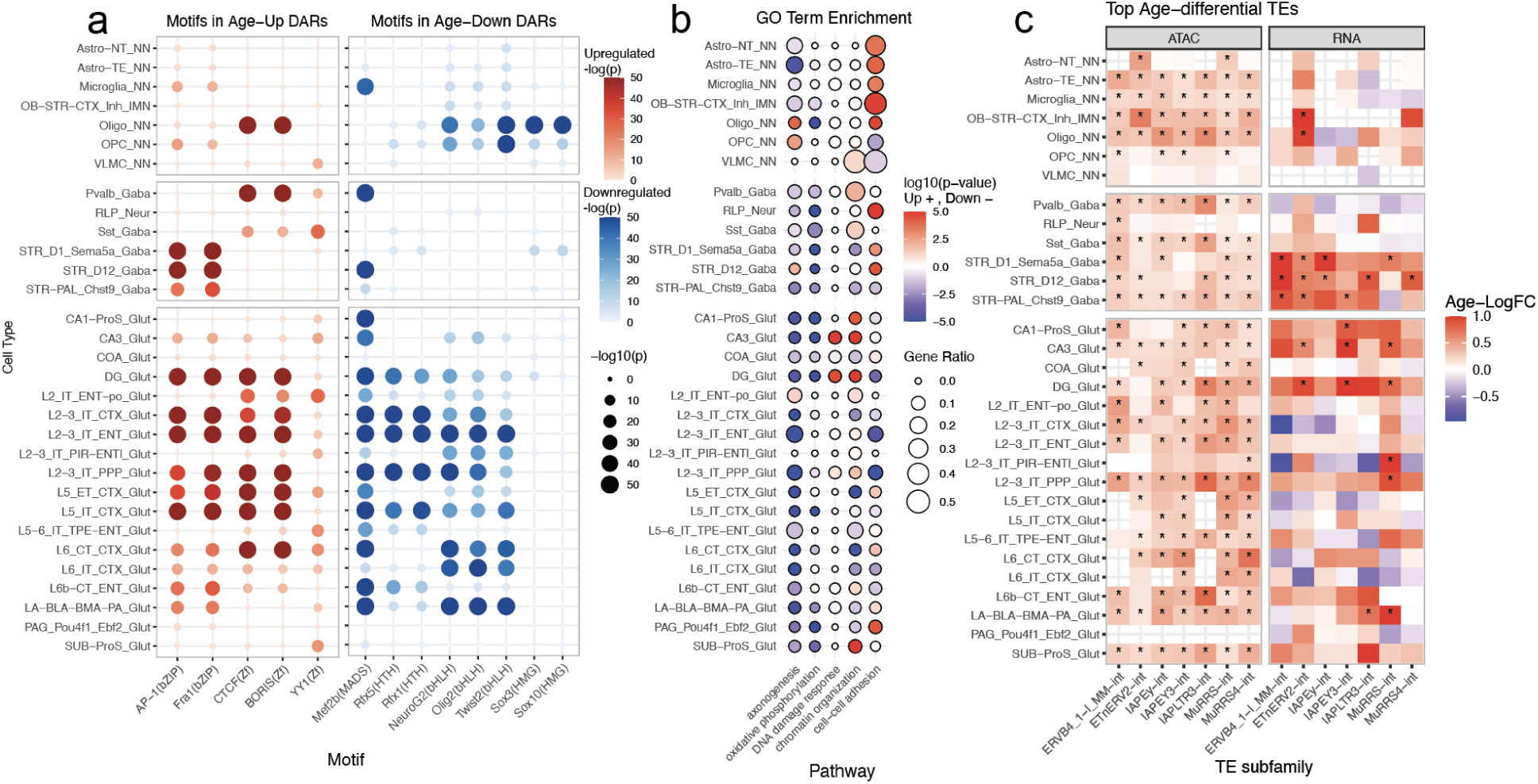
Top aging-associated regulatory and transcriptional changes across cell types. (A) Motif enrichment analysis of age-associated differentially accessible regions (age-DARs), showing top enriched transcription factor motifs across cell types. (B) Gene Ontology (GO) pathway enrichment analysis of age-associated differentially expressed genes (age-DEGs), highlighting recurrent biological processes altered with aging. (C) Top differentially accessible transposable element (TE) subfamilies across aging, ranked by frequency and magnitude of accessibility changes.

**Figure S13.**
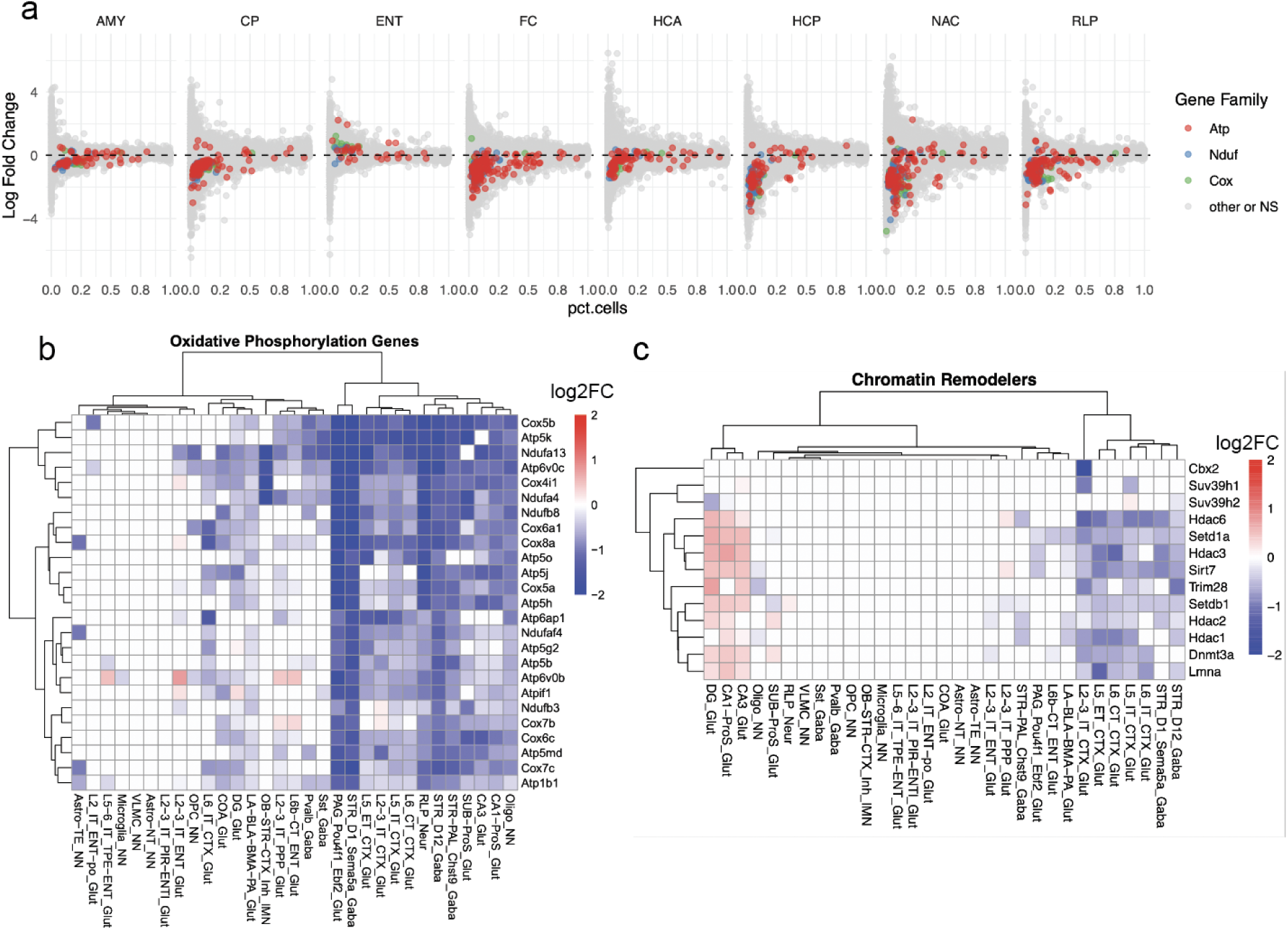
Aging-associated downregulation of oxidative phosphorylation and heterochromatin maintenance genes. (A) MA plots highlighting age-associated downregulated genes (18mo vs. 2mo) related to oxidative phosphorylation across brain regions. (B) Heatmap showing log_2_ fold change (log_2_FC) in expression of top oxidative phosphorylation-related genes across major cell types. (C) Heatmap showing log_2_FC in expression of genes involved in heterochromatin maintenance, indicating widespread transcriptional decline of chromatin regulators with aging.

## Methods

### Mouse Brain Tissue Dissection and Nuclei Isolation

All experimental procedures using live animals were performed at the Salk Institute, and were approved by the Animal Care and Use Committee (IACUC). The Salk Institute has approved Assurance from NIH Office of Protection from Research Risks (D16-00328). C57BL/6J male and female mice were purchased from the Jackson Laboratory at 7, 38 and 63 weeks of age and maintained in the Salk animal barrier facility on 12-h dark–light cycles with food ad libitum until brain extraction (housing conditions: temperature of 21–23 °C, relative humidity of 61–63%). For synchronizing the estrous cycle in females ahead of brain extraction, females were exposed to male bedding for three days such that they would be in di-estrous at time of dissection. All animals were dissected between 1 and 5 pm to ensure similarity of circadian rhythm. Brains were extracted at 2, 9 or 18 months of age, sliced coronally at 600 µm and regions dissected in an ice-cold dissection buffer as previously described^56^ according to the Allen Brain Reference Atlas CCF (version 3)^56^(CCFv3). Dissected tissue corresponding to the same region (across the anterior posterior axis) from each animal was collected across brain slabs and stored at −80 °C. For nuclei isolation, region pools from 3–14 animals were processed for each of 2–3 biological replicas per region. Comprehensive brain dissection metadata are provided in Table S1. No statistical methods were used to predetermine sample sizes. We empirically determined the number of animals needed to obtain sufficient nuclei from each brain region to run single nuclei sequencing experiments. We use two to three biological replicas for all single-cell epigenomic experiments to achieve minimum reproducibility for large-scale projects. Blinding and randomization was not performed during handling of the tissue samples. Additionally, all dissected regions were previously digitally registered into CCFv3 using ITK-SNAP^56^ (v.4.0.0) at 25 μm resolution. Nuclei isolation was achieved following established protocols as previously described^56^.

### Single-nucleus ATAC-seq Data Generation

Nuclei were pelleted with a swinging bucket centrifuge (500g, 5 min, 4°C; 5920R, Eppendorf). Nuclei pellets were resuspended in 1 ml nuclei permeabilization buffer (1mM DTT (D9779, Sigma), 0.2% IGEPAL-CA630 (Sigma), 1X cOmplete EDTA-free protease inhibitor (05056489001, Roche) and 5% BSA (A7906, Sigma) in PBS) and pelleted again (500g, 5 min, 4°C; 5920R, Eppendorf). Nuclei were resuspended in 500 uL of tagmentation buffer (36.3 mM Tris-acetate, pH 7.8 (BP-152, Thermo Fisher Scientific), 72.6mM Potassium-acetate (P5708, Sigma), 12.1 mM Magnesium-acetate (M2545, Sigma), 17.6% DMF (DX1730, EMD Millipore) in molecular biology-grade water) and counted using a hemocytometer. Concentration was adjusted to 2000 nuclei/uL, and 2000 nuclei were dispensed into each well of a 96-well plate. For tagmentation, 1 uL of barcoded Tn5 transposomes were added using a BenchSmart 96 (Mettler Toledo, RRID:SCR_018093; table S26), mixed five times, and incubated for 60 min at 37°C with 500 rpm shaking. To inhibit the Tn5 reaction, 10 uL of 40 mM EDTA was added to each well with a BenchSmart 96 and mixed 5 times, then incubated at 37°C for 15 min with 500 rpm shaking. Next, added 20 μl 2x sort buffer (2 % BSA, 2 mM EDTA in PBS), mix 5x and combine in a reagent reservoir. If processing two or more samples per day, tagmentation was performed with different sets of barcodes in separate 96 well plates. After tagmentation nuclei from individual plates were pooled together, stained nuclei suspension with 3 μM Draq7 (1:100, 7406, Cell Signalling). Sort 20 nuclei per well into eight 96-well plates (total of 768 wells, 15,360 nuclei per sample) containing 10.5 ml EB (25 pmol primer i7, 25 pmol primer i5, 200 ng BSA (Sigma). Preparation of sort plates and all downstream pipetting steps were performed on a Biomek i7 Automated Workstation.

After addition of 1 uL 0.2%SDS (15553035, Thermo Fisher Scientific) to each well, samples were incubated at 55°C for 7 min with 500 rpm shaking. Next, 1 uL 12.5%Triton-X () was added to each well to quench the SDS. Next, 12.5 ml NEBNext High-Fidelity 2X~ PCR Master Mix (M0541, NEB) were added and samples were PCR-amplified (72°C 5 min, 98°C 30 s, (98°C 10 s, 63°C 30 s, 72°C 60 s) X 12 cycles, held at 12°C). After PCR, contents in all wells were combined. Libraries were purified according to the MinElute PCR Purification Kit manual (Qiagen) using a vacuum manifold (QIAvac 24 plus, Qiagen) and size selection was performed with SPRI Beads (Beckmann Coulter, 0.55x and 1.5x). Libraries were purified one more time with SPRI Beads (Beckmann Coulter, 1.5x). Libraries were quantified using a Qubit dsDNA HS Assay Kit (Invitrogen) and the nucleosomal pattern was verified using a Tapestation (High Sensitivity D1000, Agilent). Libraries were sequenced on a NextSeq500 (Illumina) and NovaSeq6000 (Illumina) using custom sequencing primers with following read lengths: 50 + 10 + 12 + 50 (Read1 + Index1 + Index2 + Read2).

### 10X Multiome (snATAC + snRNA) Data Generation

Nuclei were pelleted with a swinging bucket centrifuge (1000g, 10 min, 4°C; 5920R, Eppendorf). Nuclei pellets were resuspended in sort buffer (1 % BSA, 2uM 7-AAD, 1x cOmplete EDTA-free protease inhibitor and 1U/uL RNasin in PBS) and stained 10 mins on ice. Nuclei were sorted, and 120,000 nuclei were collected in 90 μL of collection buffer (5 % BSA and 5U/uL RNasin). Nuclei were centrifuged (500 × g for 5 min at 4°C), and resuspended in nuclei permeabilization buffer (1mM DTT (D9779, Sigma), 0.2% IGEPAL-CA630 (Sigma), 1X cOmplete EDTA-free protease inhibitor (05056489001, Roche) and 5% BSA (A7906, Sigma) in PBS) incubated on ice for 1 min. Added 900 uL wash buffer and and pelleted again (500g, 5 min, 4°C; 5920R, Eppendorf). The nuclei were resuspended in 10 μL of 1X Nuclei Buffer and counted using a hemocytometer. A total of 18,000 nuclei were loaded per tagmentation reaction loaded onto a Chromium Controller (10x Genomics) for downstream 10X Multiome ATAC and gene expression workflows. Final library concentration was assessed by Qubit dsDNA HS Assay Kit (Invitrogen) and fragment size was checked using Tapestation High Sensitivity D1000 (Agilent) to ensure that proper fragment sizes. Libraries were sequenced using the NextSeq500 and a NovaSeq6000 (Illumina).

### snATAC-seq Data Processing

Reads were aligned to the mm10 reference genome using bowtie2 (v2.3)^57^ with default parameters and cell barcodes were added as a BX tag in the bam file. Only primary alignments were kept. Then we removed duplicated read pairs with Picard. Only proper read pairs with insert size less than 2000 were kept for further analysis. From the filtered BAM files, we processed the data using SnapATAC2 (v 2.5.1)^20^, following the official SnapATAC2 documentation. Barcode processing and fragment generation were performed, and doublets were removed using Scrublet^58^. Nuclei with fewer than 1,000 fragments, more than 100,000 fragments, or a TSS enrichment score below 5 were discarded. For female 10X Multiome samples, we processed the CellRanger output directly using SnapATAC2, ensuring consistency between modalities. Fragment files were imported, and doublets were identified and removed using SnapATAC2’s built-in method. TSS enrichment scores were computed using SnapATAC2’s function, and cells with fewer than 1,000 fragments, more than 15,000 fragments, or a TSS enrichment score below 5 were removed. Read counting and clustering were performed using a cell-by-bin matrix at 500-bp resolution, which was binarized for clustering. Bins overlapping the ENCODE^59^ blacklist (mm10) were removed to eliminate problematic regions. Read coverage normalization was performed using a log10(count +1) transformation followed by Z-score scaling, and bins with extreme Z-scores greater than 2 were filtered out. Feature selection was performed by retaining the top 700,000 most variable features. For dimensionality reduction, the Jaccard Index was computed, followed by diffusion maps for embedding. The top 20 eigenvectors were selected based on variance explained, and 20 nearest neighbors were computed per nucleus. The Leiden algorithm was applied for clustering, and clusters were annotated using known marker genes. Batch effects between male and female samples were corrected using Harmony^21^ prior to clustering. Low-quality clusters, including those with low TSS enrichment or suspected doublets, were removed from downstream analysis.

### snRNA-seq Data Processing

Single-nucleus RNA-seq (snRNA-seq) data were generated using the 10X Genomics Multiome platform and processed with Cellranger-arc v2.0.2 to generate raw gene expression matrices^60^. Doublets were identified and removed using DoubletFinder from the Cell Ranger output for each sample^61^. Downstream analysis was performed using Seurat v5 in R. Individual samples were then merged into a single Seurat object. Quality control was applied to remove low-quality nuclei, filtering out cells with fewer than 500 or more than 6000 detected genes, and cells with >10% mitochondrial reads. Normalization was performed using the SCTransform (SCT) workflow, which models technical noise while preserving biological variation. Dimensionality reduction was carried out with PCA, and the top 30 components were used to compute a shared nearest neighbor (SNN) graph for clustering using the Leiden algorithm. UMAP was used for visualization. Following initial clustering, further manual filtering was applied to remove remaining doublets and low-quality subclusters. Specifically, subclusters containing >50% cells with low UMI counts or high doublet scores were excluded.

### Cell Type Annotation via Label Transfer

Cell type annotation was performed using label transfer from a reference dataset generated by the Allen Brain Institute (10X V3)^23^. A subsampled version of this dataset, retaining 500 cells per cluster, was used as a reference. Subclass labels were transferred using Seurat’s label transfer framework with rpca. Labels were transferred for gabaergic, glutamatergic, and non-neuronal cell types separately. Each subcluster of our scRNA-seq data was assigned a label if more than 80% of the cells within that subcluster were consistently mapped to a single reference label. For ambiguous clusters where label transfer results were inconsistent, manual annotation was performed using marker gene expression profiles to ensure accurate classification. This approach resulted in the identification of 80 distinct cell types in the snRNA-seq dataset. Given that the data was 10X multiome, these annotations were also available for the corresponding ATAC-seq data. We extended the labels to the male scATAC-seq data via our joint clustering of the data. Subclusters were assigned labels based on whether the majority of labelled nuclei—previously labeled through RNA-based annotation—mapped to a specific cell type. This ensured consistency in cell type annotation across both RNA and ATAC datasets.

### MERFISH Gene Panel Design

To investigate cell-type-specific and age-associated transcriptional changes in the mouse brain, we designed a customized MERFISH panel consisting of 500 genes. Gene selection was informed by single-cell and spatial transcriptomic datasets, focusing on both cellular identity and biological processes implicated in aging. The panel includes 328 genes selected as cell type markers, capturing major neuronal and glial subtypes across brain regions. Marker genes were derived from comprehensive single-cell RNA-seq atlases^19,62^ and a MERFISH-based spatial atlas of the aging mouse brain^16^. From the latter, we prioritized genes with the highest variance across cells to ensure inclusion of the most informative spatial markers. To increase resolution for early neurogenic populations, we also included a focused subset of genes enriched in neural progenitor populations, such as radial glia-like cells and dentate gyrus progenitors. The remaining 172 genes were selected for their relevance to aging biology. These include genes differentially expressed with age in previous datasets, as well as genes involved in pathways such as chromatin remodeling, inflammation, oxidative stress, and circadian regulation. A subset of 24 genes was included based on prior evidence linking clock dysfunction to aging phenotypes^63^. MERFISH panel genes and their corresponding sources can be found in Table S2.

### MERFISH tissue preparation and imaging

Brain slabs were embedded in OCT, rapidly frozen in isopentane and dry ice, and stored at −80 °C until ready for slicing. Coronal sections (12 μm thick) were collected from OCT-embedded tissue using a Leica CM1950 cryostat at −20 °C. Sections were immediately fixed in 4% paraformaldehyde (pre-warmed to 37 °C) for 30 minutes, then permeabilized in 70% ethanol according to the manufacturer’s instructions. Sample preparation, including probe hybridization and gel embedding, was performed using the Vizgen sample preparation kit (10400012) following the provided protocol. Imaging was conducted with a MERSCOPE 500 Gene Imaging kit (Vizgen, 10400006) on a MERSCOPE system (Vizgen).

### MERFISH data preprocessing

Following the data processing performed by the Vizgen MERSCOPE, the MERFISH data was first re-segmented using CellPose^64^ (v2.1.1). Using the segmentation pipeline defined by the CellSegmentation class of the mftools package, the CellPose‘cyto-2’ model was used to perform a 2D segmentation on the 3rd z-slice of the DAPI stain images. These new segmentation masks were then used to regenerate the cell by gene matrix for downstream analysis. The MERFISH datasets were initially filtered to remove cells with less than 5 genes detected and to remove genes found in less than 3 cells. The datasets across all age groups were concatenated and processed using common Scanpy^65^ (v1.10.0) functions in Python (v3.10.14). The datasets were first normalized to have the same total transcripts per cell and then log scaled, followed by principal component analysis (PCA). The nearest neighbors distance matrix was calculated with n_neighbors=30, then leiden clustering, PAGA, and UMAP were performed with default parameters. The combined object with 2 month, 9 month, and 18 month samples was then batch corrected using Scanpy’s ComBat function. To perform a preliminary annotation of the processed MERFISH samples, the Allen Brain Cell Atlas 10Xv3 single-cell mouse brain dataset was used as a reference dataset^23^. The subclass level annotation was applied to the single-cell reference data, and this label was then transferred to the MERFISH data using the label_clusters function from the mftools package. The label_clusters function identifies the common genes between the MERFISH and reference data, then computes the mean z-scored expression of each cluster in the MERFISH and reference single-cell datasets. To transfer labels from the reference data to MERFISH, the Pearson correlation is computed between each MERFISH and reference cluster’s expression profile, then the MERFISH cluster is assigned the label of its most correlated single-cell reference cluster. This resulted in 34 different cell type labels that were transferred to the MERFISH dataset as an initial set of annotations.

### Label transfer between snRNA and MERFISH for annotation

To assign cell type annotations to our MERFISH single-cell transcriptomic data, we performed label transfer from a matched snRNA-seq dataset using Seurat. We first subsampled the snRNA-seq reference to reduce the overrepresentation of large clusters, capping the maximum number of cells per annotated cluster at 2,000. Both the snRNA-seq and MERFISH datasets were normalized using SCTransform, and transfer anchors were computed using canonical correlation analysis (CCA) with the top 45 dimensions and all available genes (FindTransferAnchors, reduction = “cca”). Labels from the RNA-seq data (celltype_final) were transferred using TransferData, generating a predicted label and a confidence score (prediction.score.max) for each MERFISH cell. To mitigate potential noise and improve robustness, we refined the transferred labels through subclustering. Each MERFISH cluster sub-clustered at higher resolution (FindClusters, resolution = 2) following PCA-based preprocessing. For each resulting subcluster, we determined the majority predicted label based on cells with high confidence (score > 0.85). Any ambiguous subclusters were removed. For both modalities, gene expression was aggregated across cells of the same cell type using Seurat’s AggregateExpression() function with centered log-ratio (CLR) normalization. The CLR transformation was applied per cell to adjust for differences in sequencing depth and compositional effects. For the RNA-seq data, gene expression values were further normalized by gene length to account for transcript length bias, yielding RPKM-like units based on CLR-normalized counts. Pearson correlation was computed across shared genes and cell types between the RNA-seq and MERFISH aggregated matrices. The resulting gene-by-gene correlation matrix was visualized as a heatmap to assess concordance in gene expression patterns across modalities (Figure S7).

### Differential Gene Expression Analysis

Differential gene expression analysis was conducted using MAST^26^, incorporating latent variables to account for percent mitochondrial reads, percent ribosomal reads, replicate, and brain region where necessary. Statistical significance was determined using likelihood ratio tests, and multiple comparisons were adjusted using the Benjamini-Hochberg correction. Genes were considered differentially expressed if they met a log2 fold-change of 0.25 and an adjusted p-value < 0.01.

### Pathway Enrichment Analysis

Pathway enrichment analysis was performed using clusterProfiler^66^ on significantly upregulated and downregulated genes with all genes used in the DEG analysis as background (genes expressed in > 1% of cells).

### Peak Calling and Differential Accessibility Analysis

Peak calling and differential accessibility analysis were performed using SnapATAC2^20^. For peak calling, MACS3 was run on individual cell types using the snap.tl.macs3 function. Peaks were called separately for each cell type and age group using a q-value threshold of 0.01, with additional filtering against the ENCODE mm10 blacklist. To ensure robust peak detection, replicate-specific peaks were discarded. To identify differentially accessible regions (DARs), comparisons were made between 2-month-old and 18-month-old nuclei within each cell type. Peaks were filtered to include only those detected in at least 1% of cells in either age group. Differential testing was performed using snap.tl.diff_test, with a minimum log2 fold change of 0.25 and a significance threshold of adjusted p-value < 0.01. Up- and downregulated peaks were extracted and saved as BED files for downstream analysis. For visualization, aggregated peak matrices were generated for each cell type, region, and age group, normalized to reads per million (RPM).

### Motif Enrichment Analysis

Motif enrichment analysis was performed using HOMER (v 4.11)^67^ for the age-differential peaks in each cell type, with non-differential peaks as the background and default parameters.

### Overlap with histone marks

We used bedtools intersect -c to overlap all called peaks for each cell type with H3K9me3 data from ENCODE^46^. H3K9me3 data from the forebrain were re-aligned to mm10 genome using BWA^68^ without mapping quality filter (in order not to lose any reads aligning to repetitive elements), and peaks were re-called using SICER^69^ on both ChIP-seq and input libraries, as described in^17^.

### Linking Genes to cCREs via ABC score

To associate genes with putative regulatory elements, we employed the Activity-By-Contact (ABC) model, which integrates chromatin accessibility, histone modifications, and 3D chromatin interactions to predict enhancer-gene links. The ABC score was calculated following the framework described by Fulco et al.^36^, using open chromatin peaks as candidate cis-regulatory elements (cCREs).

### Gaussian Smoothing for Hotspot Identification

We used the R package smoother to perform Gaussian smoothing on the number of differentially accessible regions (DARs) (p < 0.01, |log2 fold change| > 1) within each 100 kb region of the genome. A smoothing window length of 20 bins (~2.1 Mb span) was applied to account for local clustering of accessibility changes. Regions of the genome with a high concentration of DARs within a short distance were assigned higher Gaussian smoothing scores, and hotspots were defined as bins with smoothing scores in the top 1%, corresponding to a threshold of 0.2.

### Quantifying transposable elements (TEs) accessibility

scTE^70^ version 1.0 was used to build a genome index for the alignment of reads to genes (gencode vM21) and TEs (rmsk mm10) using scTE_build. The scTE command was used to map reads from the BAM files to transposable element families, generating a cell by feature read count matrix. A pseudo-bulk count table fo TE subfamilies was generated by summing reads from cells of the same cell type, age and biological replicate for each feature. Age-differential TEs for each cell type were then identified by DeSeq2^71^ between 18-month and 2-month datasets using the likelihood ratio test.

### Quantifying transposable elements (TEs) expression

SoloTE^72^ was used to count reads at TE elements and genes directly from the Cellranger output, generating a Seurat object that contained the gene and TE count matrix. A pseudo-bulk count table for both genes and TEs was generated by summing reads from cells of the same cell type, age and biological replicate for each feature. Age-differential genes and TEs for each cell type were then identified by DeSeq2^71^ between 18-month and 2-month datasets using the likelihood ratio test.

## Supplementary tables

Table S1. Brain Dissection Region Annotation

Table S2. MERFISH gene panel

Table S3. ATAC Metadata

Table S4. RNA Metadata

Table S5. MERFISH Metadata

Table S6. cCREs called by celltype table

